# Transcriptomic Analyses Throughout Chili Pepper Fruit Development Reveal Novel Insights into Domestication Process

**DOI:** 10.1101/2020.10.05.326470

**Authors:** Octavio Martínez, Magda L. Arce-Rodríguez, Fernando Hernández-Godínez, Christian Escoto-Sandoval, Felipe Cervantes-Hernández, Corina Hayano-Kanashiro, José J. Ordaz-Ortiz, M. Humberto Reyes-Valdés, Fernando G. Razo-Mendivil, Ana Garcés-Claver, Neftalí Ochoa-Alejo

## Abstract

Chili pepper (*Capsicum* spp.) is both an important crop and a model for domestication studies. Here we performed a time course experiment to estimate standardized gene expression profiles across fruit development for six domesticated and four wild chili pepper ancestors. We sampled the transcriptome every 10 days, from flower to fruit at 60 Days After Anthesis (DAA), and found that the mean standardized expression profile for domesticated and wild accessions significantly differed. The mean standardized expression was higher and peaked earlier for domesticated vs. wild genotypes, particularly for genes involved in the cell cycle that ultimately control fruit size. We postulate that these gene expression changes are driven by selection pressures during domestication and show a robust network of cell cycle genes with a time-shift in expression which explains some of the differences between domesticated and wild phenotypes.

## INTRODUCTION

Chili peppers of genus *Capsicum* and Solanaceae family are native to the American continent. Of the approximately 30 chili pepper species, five have been domesticated: *C. annuum* L., *C. frutescens* L., *C. baccatum* L., *C. chinense* Jacq. and *C. pubescens* Ruiz & Pav. (Pickersgill, 1971). Among these species, *C. annuum* is the most important worldwide as a vegetable and spice crop, and production of this type of pepper has been steadily increasing both in terms of area harvested and yield (Jarret et al., 2019). In addition to its economic importance, chili peppers are a source of antioxidants such as flavonoids, phenolic acids, carotenoids and vitamins (Badia et al., 2017; Cervantes-Hernández et al., 2019), as well as a plant model to study the genetic and biochemical basis for synthesis of these compounds (Gómez-García and Ochoa-Alejo, 2013; Gómez-García and Ochoa-Alejo, 2016; Martínez-López et al., 2014). Capsaicinoids, which are synthesized only in *Capsicum* species, impart pungency to chili peppers and are a focus of active research (Arce-Rodríguez and Ochoa-Alejo, 2017; Tanaka et al., 2017; Fayos et al., 2019).

Chili peppers were domesticated from an ancestral variety, *Capsicum annuum* L. var. *glabriusculum*, locally known as “piquín” or “chiltepín” (Hayano-Kanashiro et al., 2016) in northeastern Mexico and/or central-east Mexico (Kraft et al., 2014). The oldest chili pepper macroremains date to around the time of first cultivation or domestication in the mid-Holocene, 9,000-7,000 BP (Kraft et al., 2014; McClung de Tapia, 1992). The exact time of chili pepper domestication is a subject of debate (Pickersgill, 2016), as starch microfossils of domesticated *Capsicum* dating from 6,000 BP have been found at seven sites (Perry et al., 2007), and there is evidence indicating that the fruit size of domesticated genotypes has increased considerably in the last 1,500-1,000 years BP (Pickersgill, 2016). Larger fruit size in domesticated compared with wild ancestors is part of the “domestication syndrome” (Doebley et al., 2006).

Domestication, which involves breeding and selection of wild ancestral forms to modify phenotypes for human use, is not only a key achievement of modern civilization (Zeder, 2015), but also provides a unique opportunity to identify the genetic basis of adaptation (Ross-Ibarra et al., 2007). Examples of studies of plant domestication include maize (Doebley et al., 1990; Tian et al., 2009; Studer et al., 2011; Hufford et al., 2012), common bean (Bellucci et al., 2014; Singh et al., 2018), tomato (Lippman and Tanksley, 2001; Müller et al., 2016; Sauvage et al., 2017; Razifard et al., 2020) and *Capsicum* (Hernández-Verdugo et al., 2001; Paran and Van Der Knaap, 2007; Carvalho et al., 2014; Taitano et al., 2019).

The *Capsicum* genome (∼3.5 Gb) has been sequenced and annotated (Qin et al., 2014; Kim et al., 2014); currently there are 9 genomic assemblies available in the NCBI (https://www.ncbi.nlm.nih.gov/genome/?term=Capsicum) and further sequencing of different genotypes has been reported (Ahn et al., 2016; Hulse-Kemp et al., 2018). In particular, Qin et al. (2014) provided insights to evaluate the adaptive landscape of cultivated peppers (Albert and Chang, 2014) and reported a set of 511 genes that have a strong genomic domestication footprint.

To study the divergence caused by domestication in gene expression profiles during chili pepper fruit development, we examined fruit transcriptomes of six domesticated and four wild accessions by RNA-Seq every 10 days from anthesis until fruiting at 60 DAA. Our data show that there are significant differences in the mean expression profiles of domesticated and wild accessions that affected a set of interrelated biological processes, particularly the cell cycle. We postulate that such differences in expression profiles could partially explain the large difference in fruit size between domesticated and wild chili pepper varieties.

## RESULTS

We constructed and analyzed RNA-Seq libraries from developing fruits at seven time points (0, 10, 20, 30, 40, 50 and 60 DAA) from 10 accessions (6 Domesticated (D) and 4 Wild (W)). Table 1 shows the key, names and accession type.

**Table 1.** Chili pepper accessions used in this study

| Domesticated (D) |  | Wild (W) |  |
| --- | --- | --- | --- |
| Key | Name | Key | Name |
| AS | Ancho San Luis | CO | Piquín Coahuila |
| CW | California Wonder | QU | Piquín Querétaro * |
| CM | Criollo de Morelos 334 * | SR | Piquín Sonora Red |
| JE | Jalapeño Espinalteco | SY | Piquín Sonora Yellow |
| ST | Serrano Tampiqueño |  |  |
| ZU | Zunla * |  |  |
| * - Genome available for CM (Kim et al., 2014), QU and ZU (Qin et al., 2014). |  |  |  |

A total of 22,427 genes, representing approximately 64% of the genes annotated in the *Capsicum* genome, were consistently expressed during fruit development. Data normalization is a crucial step in gene expression studies (Wu et al., 2019). For our analyses we used the Standardized Expression Profile (SEP), which is a 7-dimensional vector formed by the estimated means of expression at each time point (0, 10, 20, 30, 40, 50 and 60 DAA) that has a mean and standard deviation of 0 and 1, respectively (see Methods). The use of SEPs allows statistical comparisons between genes, or groups of genes, to be made independently of the relative gene expression of each gene. For each gene within each accession, we estimated SEPs and analyzed differences between D and W genotypes.

### Domesticated (D) and Wild (W) Accessions Have Different Mean SEPs

To study similarities in gene expression profiles between accessions, we calculated the mean Euclidean distances between SEPs for all 22,427 genes expressed during fruit development to generate a dendrogram (Figure 1).

**Figure 1.**
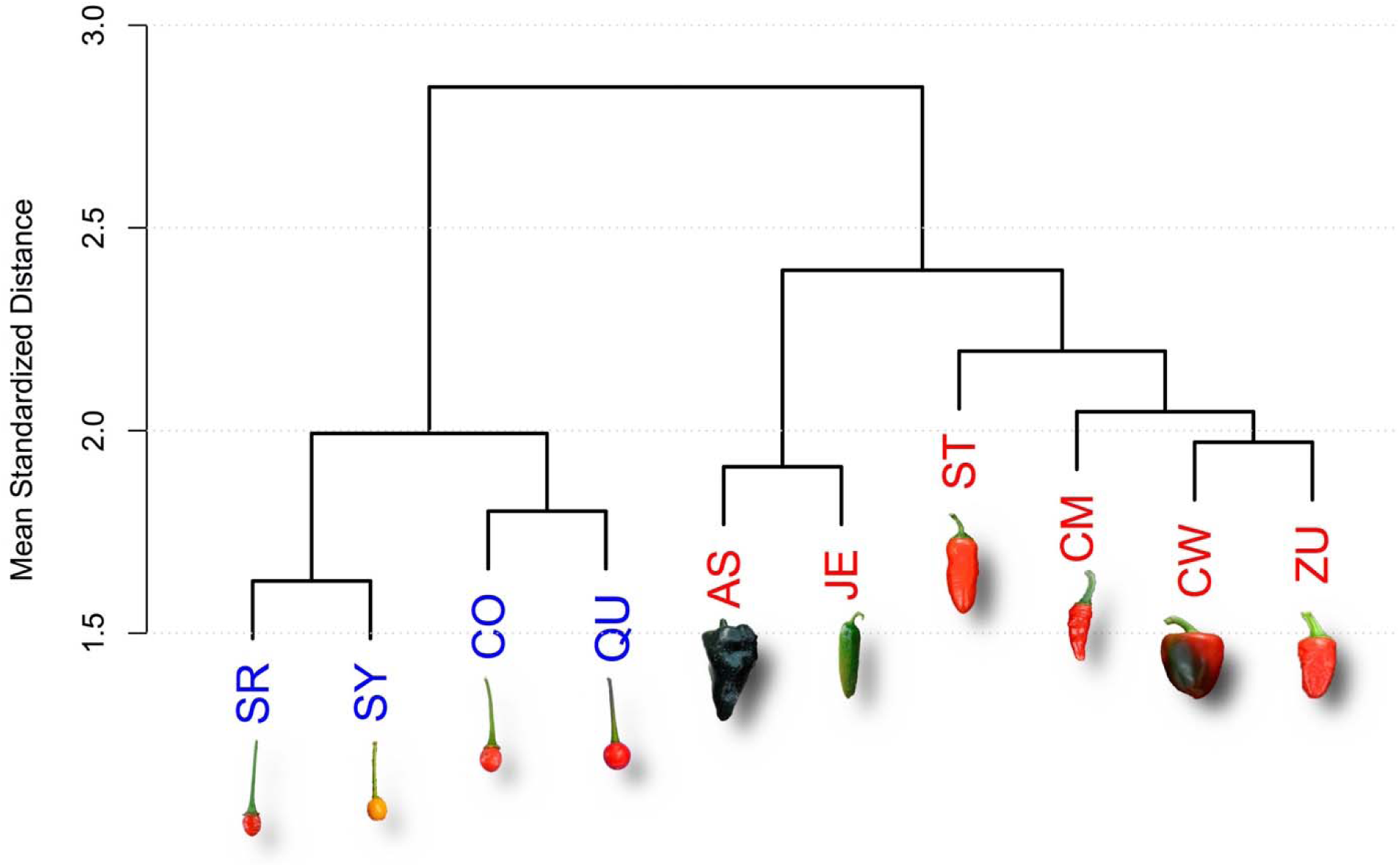
Dendrogram for Domesticated (D, red) and Wild (W, blue) Accessions. The dendrogram was obtained from the Euclidean distances between the full set of 22,427 SEPs of genes expressed during fruit development. Representative miniature photographs illustrate the fruits of the corresponding accessions. See Table 1 for accession names. Photographs of fruits are not at the same scale.

The D and W accessions form two clearly segregated groups with a mean normalized distance of 2.85 on the Y-axis (Figure 1). The four W accessions (in blue) form a cluster at a mean distance of 2, whereas the six D accessions form a cluster at a mean normalized distance of approximately 2.4 (Figure 1, and Supplemental S-3).

To perform statistical analyses of a gene, or sets of genes, we considered contrasts between two groups of accessions, 6 D (AS, CW, JE, ST and ZU in Table 1) and 4 W (CO, QU, SR and SY in Table 1). In all cases, the null hypothesis was that at each time point the mean expression of the D and W groups was equal, whereas the alternative was that these parameters differed. Variation within the D and W groups was considered as a statistical error (unexplained variation) and a t-test was used to obtain Confidence Intervals (CI) for the means and to evaluate significance at each of the 7 time points sampled. We determined the mean SEPs for different gene groups in the D and W accessions (Figure 2).

**Figure 2.**
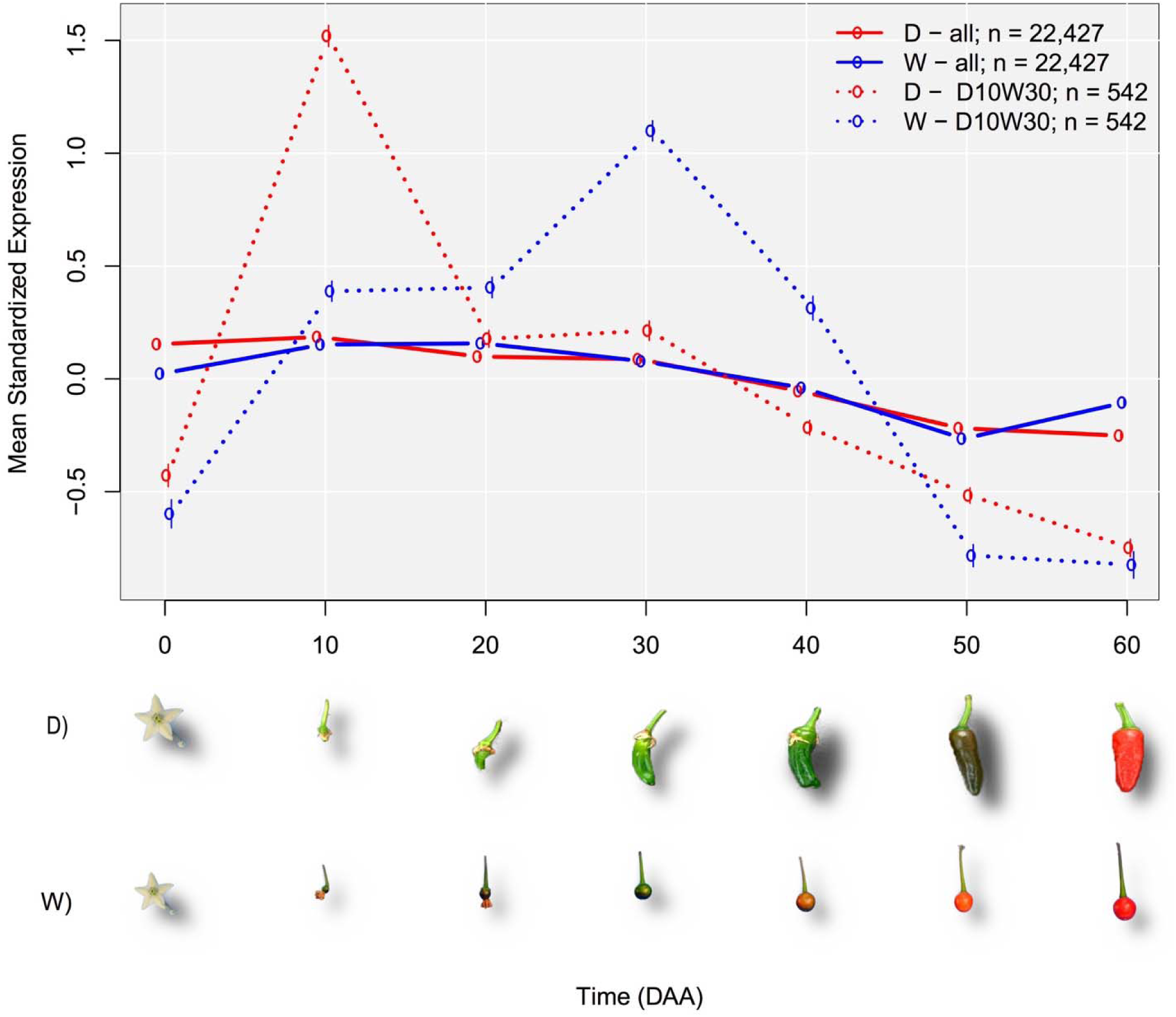
Mean SEP (Standardized Expression Profile) for Groups of Genes in Domesticated (D) and Wild (W) Accessions. Continuous colored lines (red and blue for domesticated (D) and Wild (W) respectively), link the means of standardized gene expression at each time point for the complete set of expressed genes (n=22,427). Dashed lines link the means of standardized gene expression at each time point for the n=542 genes that presented the maximum expression at 10 DAA in D while the maximum expression was reached at 30 DAA in W. These 542 genes form the group “D10W30” (see text). Representative miniature photographs illustrate the approximate fruit development of D and W accessions at each time point. Photographs of fruits are not at the same scale.

The mean for the D and W groups differed significantly (continuous line in Figure 2). At the mature flower state (0 DAA), the standardized mean expression for D was much higher than for W, implying that the average transcription activity in this state is substantially larger for the D genotypes. In the interval between 0 and 10 DAA, the mean standardized expression increased for both groups, although the rate of increase was higher for D. At 10 DAA, the mean expression for D reached a peak value, but for W the increase continued, although at a slower rate, to peak at 20 DAA. From the peak at 10 DAA, the mean expression for D decreased, at different rates, and was lower at all subsequent time points. The lowest value was seen at 60 DAA. In contrast, decreases in the mean expression for W began later, occurring from 20 up to 50 DAA, and reached a minimum of −0.27, which is smaller than the minimum for the D group, −0.25, seen at 60 DAA. The more relevant differences in the mean expression profiles between D and W were seen during the intervals 10 to 20 and 50 to 60 DAA, when the trend was inverted such that D was decreasing while W was increasing. On the other hand, less marked differences between D and W were seen between 30 and 50 DAA when the mean standardized expression decreased nearly in parallel for both groups. The average of the time at which the maximum expression was reached in each group was five days earlier for D than W. All observed differences were significant (see Supplemental S-4).

Differences in SEP of individual genes varied between D and W. A total of 463 genes, representing approximately 2.06% of the total, show significant differences between D and W with a False Discovery Rate (FDR) threshold of 0.05, which for the individual tests corresponds to a *P* value < 0.000002. The differences in expression profiles between D and W were well defined and significant; the peak of mean expression for D occurred at 10 DAA, while the peak for W occurred later, at 30 DAA. The average time of maximum expression was 11.06 DAA for D and 28.33 DAA for W, or a difference of −17.27 DAA. Of the 463 selected genes, 36 (36/463 ≈ 0.08 or 8%) are transcription factors (TFs). This percentage is higher than that for TFs annotated in the *Capsicum* genome (1,859/34,986 ≈ 0.05 or 5%). A list and description of the 463 selected genes and details of statistical analyses are presented in the Supplemental SG and S-4, respectively.

We established that the main differences in SEPs between D and W were due to a set of 542 genes which presented the expression peak at 10 DAA in D, while the expression peak was at 30 DAA in the W accessions, naming such groups of genes as D10W30 (dashed line in Figure 2).

The results of this experiment showed differences in expression profiles between D and W at the level of whole gene sets, groups of particular genes, and individual genes (see Supplemental S-4). Taking these findings together, we can thus conclude that there are relevant differences in expression profiles between domesticated and wild varieties of chili peppers during fruit development.

We also found that gene expression diversity, expressed as the coefficient of variation of gene expression, is significantly (*P* = 0.002) smaller in D than W accessions, corroborating the findings presented by Liu et al. (2019) for different species of plants and animals.

### Differences in Expression of Genes Related to Cell Reproduction Appear Earlier and are Larger in Domesticated than Wild Genotypes

Based on the evidence that mean SEPs differ between the D and W accessions, we investigated differences in expression profiles in groups of genes related to particular biological processes. We first examined the mean SEPs of a group of 1,125 genes associated with cell reproduction (Supplemental S-4.1).

The mean expression value for 235 genes that are directly annotated in the cell cycle–but not in other cell reproduction processes–was significantly higher and occurred earlier for D compared to W, as evidenced by the peak of 0.3 standardized units at 10 DAA for D and 0.2 standardized units 30 DAA for W (Supplemental S-4.1). Similarly, the mean expression for 69 kinesins or kinesin-related proteins among the 1,125 genes associated with cell reproduction exhibited a differential expression peak at 10 DAA for D accessions, but for W accessions the peak was later at 30 DAA (Supplemental S-4.1).

Thus, changes in expression of genes associated with cell reproduction were significantly larger and occurred earlier for D relative to W accessions, not only for the full set of genes, but also for particular bioprocesses and gene families (Supplemental S-4, S-5 and S-6).

### Biological Processes Enriched in Genes That Are Expressed Earlier in Domesticated Genotypes

The results presented before indicate that SEPs in D and W accessions undoubtedly differ (Figure 2), and genes for which expression peaks at 10 DAA for D but at 30 DAA for W (denoted here as ‘D10W30’) play an important role in cell reproduction. To validate and expand our study, we considered the D10W30 expression pattern in a Gene Ontology enrichment analysis (for details see Supplemental S-4.2, S-5 and SG).

A total of 86 biological processes (BPs) were significantly enriched (FDR=0.05; P<0.0015) in the D10W30 set, with a median odds ratio of 9.5. As such, these genes were much more abundant in these BPs than would be expected by chance. Apart from the abovementioned BPs related to cell reproduction, 43 of the enriched BPs, or 50% of the total, are involved in either positive or negative regulation of various biological processes. Of these, 4 (5%) are related to cellular component organization or biogenesis, 3 are associated with cellular component assembly, and another 3 play roles in organelle organization or fission. The general bioprocess “cellular process” (GO:0009987) is also highly enriched in the D10W30 gene set, with an odds estimate of 2.25 and a highly significant *P*-value of 2.76 ⨯ 10^−8^.

These results show that genes having the pattern D10W30 are over-represented in important BPs, which in turn implies that expression of such BPs occurs earlier and at higher levels in D compared to W genotypes.

Interesting examples of D10W30 genes involved in cell reproduction are a high mobility group B protein 6 (XP_016555757.1), a MYB-related protein 3R-1 (XP_016537977.1) and the kinetochore protein NDC80 (XP_016539151.1); see Supplemental S-4.2 for SEP plots. The gene encoding the “high mobility group B protein 6”, is a WRKY transcription factor involved in the nucleosome/chromatin assembly that was annotated in 12 of the 86 abovementioned BPs, particularly cell reproduction BP. The gene encoding the transcription factor “MYB-related protein 3R-1” was included in 6 of the 86 enriched BPs and is mainly related to cellular, chromosome and organelle organization. The “kinetochore protein NDC80” is part of the multiprotein kinetochore complexes that couple eukaryotic chromosomes to the mitotic spindle to ensure proper chromosome segregation. NDC80 is part of the outer kinetochore and forms a heterotetramer with proteins NUF2, SPC25 and SPC24 (Santaguida and Musacchio, 2009; D’Archivio and Wickstead, 2017). Interestingly, the genes encoding NUF2 and SPC25 also exhibit the D10W30 expression pattern. NDC80 is conspicuously present in 74 of the 86 enriched BPs.

### A Network of Cell Cycle Genes with D10W30 Expression Pattern

The availability of genome-wide gene expression technologies, as RNA-Seq, make it possible to identify gene interactions and represent them as gene networks (Filkov et al., 2005). In functional genomics it is axiomatic that genes with highly similar expression profiles are likely to be regulated via the same mechanisms, and this hypothesis is the basis for the discovery of regulatory networks (Allocco et al., 2004). To show the concerted co-variation through time of cell cycle genes expression during fruit development, we estimated a gene network comprising six structural genes and eight TF, candidates to be regulating three of the six structural genes. Figure 3 presents the estimated gene network, while Table 2 gives the descriptions of the genes involved.

**Table 2.** Identifiers and descriptions for genes in the network of Figure 3A

|  |  |  |  |
| --- | --- | --- | --- |
| **Figure 3A legend** | **Protein id** | **Description** | **Ortholog** |
| (1) - MAP/ASE1 (1). | XP_016564755.1 | 65-kDa microtubule-associated protein 3 | AT5G51600 |
| (2) - DUF566. | XP_016538322.1 | ENDOSPERM DEFECTIVE 1 (DUF566) | AT2G44190 |
| (3) - Kinesin 3, Isoform X3. | XP_016541615.1 | Kinesin 3 isoform X3 | AT4G21270 |
| (4) - MAP/ASE 1 (2). | XP_016575449.1 | 65-kDa microtubule-associated protein 3 isoform X1 | AT5G51600 |
| (5) - POLENLESS 3. | XP_016577799.1 | Protein POLLENLESS 3 | AT4G20900 |
| (6) - Shugoshin. | XP_016548908.1 | Shugoshin-1; chromosome segregation | AT3G44960 |
| (7) - AP2/B3-TF (1). | XP_016568750.1 | AP2/B3-like TF family protein | AT5G42700 |
| (8) - HGM B, 6. | XP_016555757.1 | High mobility group B protein 6 | AT4G11080 |
| (9) - Cyclin S13-7. | XP_016543946.1 | G2/mitotic-specific cyclin S13-7 | AT3G11520 |
| (10) - SET domain protein 35. | XP_016547461.1 | SET domain; methyltransferase activity. | AT1G26760 |
| (11) - AP2/B3-TF (2). | XP_016575946.1 | AP2/B3-like TF family protein | AT5G58280 |
| (12) - FHA, PS1. | XP_016574880.1 | FHA domain-containing protein PS1 | AT1G34355 |
| (13) - Myb 3R-1. | XP_016537977.1 | Myb-related protein 3R-1 | AT4G32730 |
| (14) Tesmin / TSO1 CX3 3. | XP_016565918.1 | Protein tesmin/TSO1 CXC 3 | AT3G22780 |

**Figure 3.**
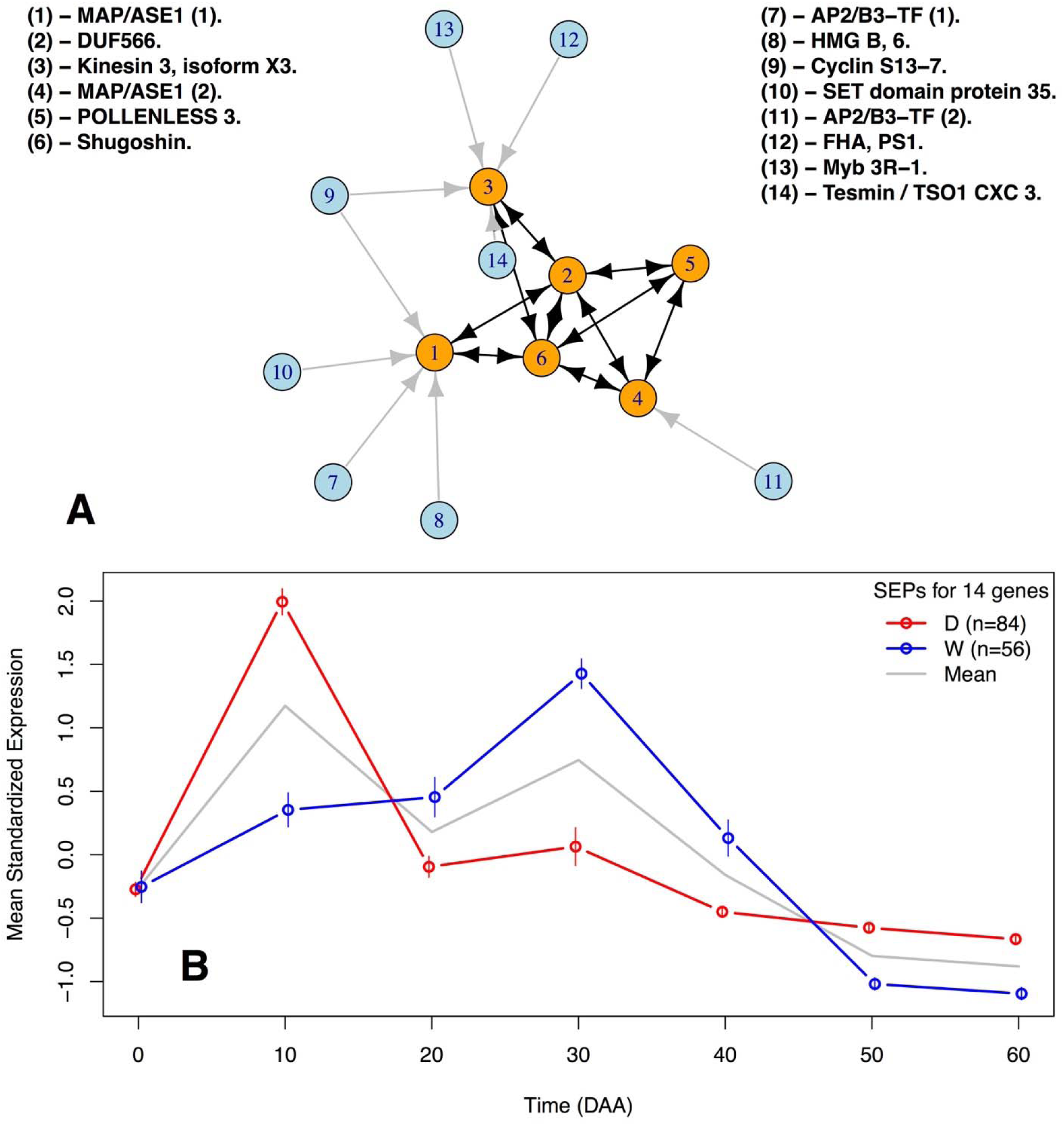
Gene Network of D10W30 Cell Cycle Genes in D and W Accessions. A - Graphic representation of the network. Orange circles represent structural genes while blue ones represent TF candidates to be regulating the genes (see Table 2 and Supplemental S-9). B - Mean standardized expression of the genes in the network in D and W accessions.

Figure 3A presents the network formed by 14 D10W30 genes, in which orange circles represent 6 structural genes involved in cell cycle, and blue circles represent TF genes candidates to be regulating 3 of the structural genes (see Figure 3A legend and Table 2 for gene descriptions). Arrows between structural genes are drawn for genes with highly significant (*P* < 0.0001) positive Pearson’s correlation coefficients (*r* > 0.96) between mean SEPs within the D and W accessions, while Pearson’s correlation coefficients of the same gene pairs between D and W accessions were small (*r* < 0.4) and not significant (*P* > 0.5). The origin and robustness of the network presented in Figure 3 A can be appreciated by observing Figure 3 B, that presents the mean SEPs for the 14 genes involved, which represent n=84 independent SEPs estimated from the 6 D accessions (red line; 84/6=14 genes) and n=56 independent SEPs estimated from the 4 W accessions (blue line; 56/4=14 genes). In Figure 3 B the 95% CI for the mean standardized expression of the genes -vertical lines at each time point, show that these 14 genes are highly correlated within D and W accessions, but have a very low correlation between the D and W groups. For details see Supplemental S-7 and S-8.

For a better appreciation of the changes that occur in the individual standardized expression of the genes included in the network of Figure 3, Figure 4 presents a grid in time and accession groups, but in this case the sizes of the circles representing genes vary in proportion to their individual standardized expression through time (rows, from 0 to 60 DAA) and sets of accessions (columns, D in the left hand side and W in the right hand side). Representative miniature photographs show the approximate fruit development stage for D and W accessions.

**Figure 4.**
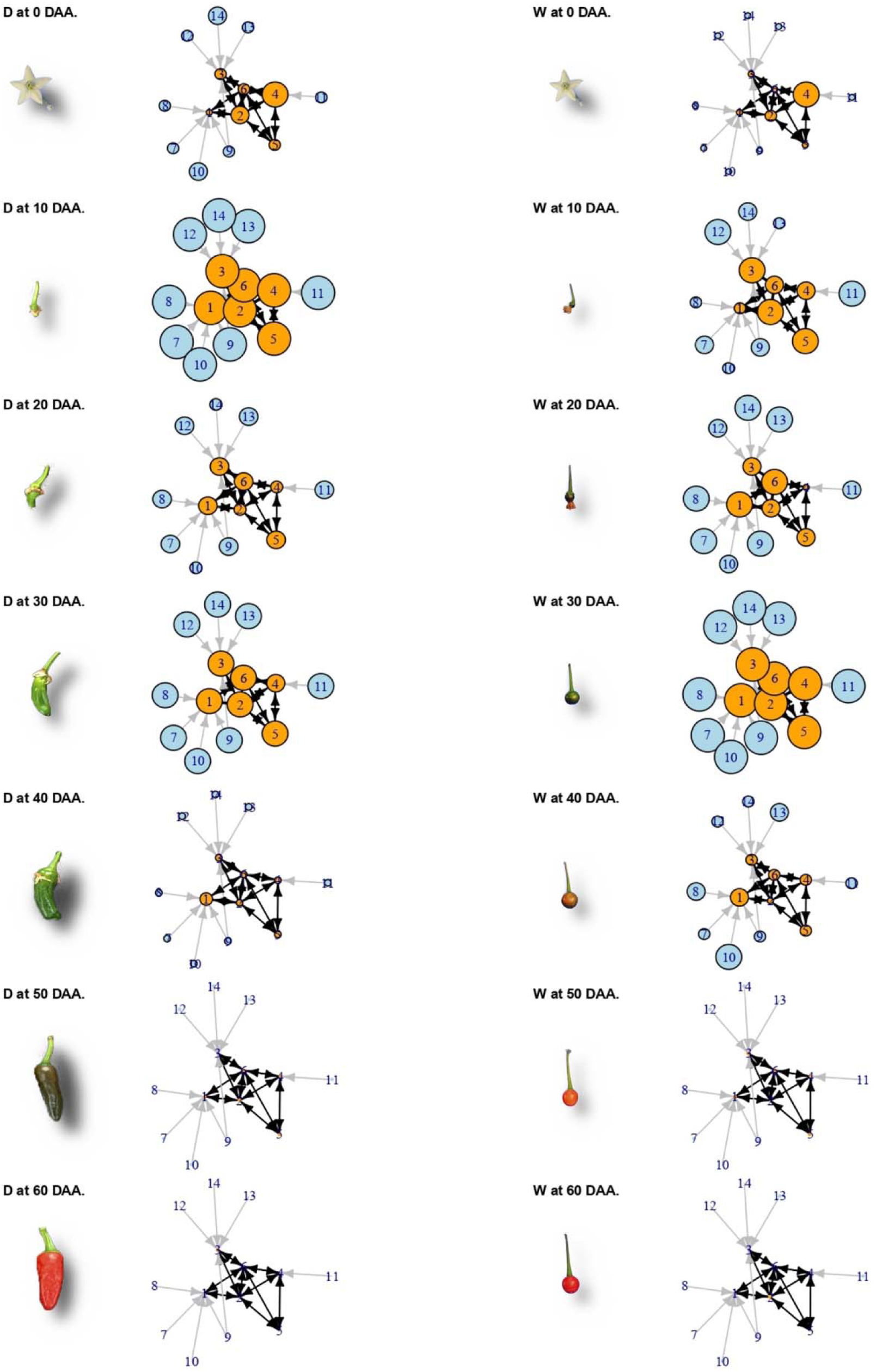
Change in Time (Rows) and Groups of Accessions (Columns) of the Network in Figure 3A. Gene expression is proportional to circle size. Photographs of fruits are not at the same scale.

In Figure 4 we appreciate that individual gene expression is highly coordinated through time, but largely differs between D and W groups of accessions. At 10 DAA in the D accessions (second row, left hand side), all 14 genes have reached their maximum expression (largest circle sizes), while such expression levels are only reached 20 days later, at 30 DAA for the W accessions (fourth row, right hand side). Thus the main cell cycle genes are well behind in maximum expression time in the W compared with the D accessions. Also, expression changes occur faster in D compared with W accessions, a fact that can be verified observing Figure 3 B, where the slop of the line linking the 0 and 10 DAA is more pronounced than the corresponding change to reach the maximum from 20 to 30 DAA in the W accessions. Departing from comparable standardized expression at the mature flower (0 DAA), individual expression rapidly diverges between D and W groups to finally converge again to basal expression levels at 50 and 60 DAA. The concerted but highly divergent expression of cell cycle genes demonstrates that domestication has tailored this process to differ between D and W genotypes, explaining in part the large differences in fruit size between these groups.

## DISCUSSION

Building on our previously described method to estimate the dynamics of the chili pepper transcriptome (Martínez-López et al., 2014), as well as our time course experiment (Spies and Ciaudo, 2015) and statistical analysis methods, in this study we generated whole gene expression profiles (SEPs) across the full course of fruit development to examine differential gene expression patterns between domesticated (D) and wild (W) varieties of chili peppers (Figure 1).

### Domestication Produced Modified Gene Expression Profiles in Chili Pepper Fruit

Domestication, as studied initially by Darwin (Darwin, 1868), is currently defined as a distinctive coevolutionary and mutualistic relationship between domesticator and domesticate, and marked a key transition in human history (Zeder, 2015). Here we propose that the differences observed between gene expression profiles in sets of domesticated (D) and wild (W) accessions (Figure 2) can be attributed to domestication.

Gene expression is an intrinsically noisy process (Swain et al., 2002); however, our experimental design systematically took into account variations between sets of fruits by examining two replicates of each accession at each time point, as well as variation within the target groups (D and W) by examining 6 and 4 genotypes for each group, respectively. Thus, the differences observed between D and W gene expression profiles can be attributed to differences in the selection history of the two groups, i.e., to domestication. The effect of domestication on gene expression patterns is also corroborated by the fact that the D and W accessions belong to well-segregated clusters in the dendrogram presented in Figure 1. Although the distance between the D and W groups is approximately 2.85, the maximum distance within the groups is only 2.4, again demonstrating that gene expression patterns during fruit development were modified by domestication.

Selection during domestication can alter molecular footprints at the genomic level. For example, in *Capsicum* Qin et al. (2014) identified 115 genomic regions containing 511 genes that show strong selective sweep signals due to domestication. The same method of searching for selective sweep signals identified candidate genes not only in plants, such as maize (Tian et al., 2009), sunflower (Chapman et al., 2008), soybean (Li et al., 2013) and Asian rice (Huang et al., 2012), but also in domesticated animals such as dogs (Pollinger et al., 2005) and cattle (Rothammer et al., 2013). Ross-Ibarra et al. (2007) reviewed methods to identify the genes responsible for adaptation to domestication and classified them in terms of the phenotype-genotype hierarchy as “top-down”, in which the method starts with a phenotype to identify candidate genes, and as “bottom-up”, in which genetic analyses are used to identify adaptive genes. Bioinformatics tools are then used to connect the selected genes to a phenotype. Our approach in this study can be considered as a hybrid between “top-down” and “bottom-up” methods in that we began with a molecular phenotype, i.e., a standardized gene expression profile (SEP), which showed that there are significant differences between D and W groups of accessions, and then examined the biological relevance of these findings.

Differences in gene expression between crops and their wild ancestors were reported in tomato (Müller et al., 2016; Dai et al., 2017), which, like *Capsicum*, belongs to the Solanaceae family, common bean (Bellucci et al., 2014; Singh et al., 2018), carrot (Rong et al., 2014), ramie (Liu et al., 2014) and cotton (Chaudhary et al., 2008). For animals, similar comparisons were done for turkey (Monson et al., 2016), trout (Christie et al., 2016) and other species (Albert et al., 2012). All of these studies reported sets of differentially expressed genes between the domesticated and wild forms, but only for tomato did the study authors use a time course experiment to evaluate deceleration of the circadian clock (Müller et al., 2016). On the other hand, a decrease in gene expression diversity in domesticated forms of a set of animals and plants was reported by Liu et al. (2019). Analysis of our data confirmed that gene expression is significantly less diverse in domesticated chili peppers compared to that seen for wild accessions.

### Biological Relevance of the Differences Between Gene Expression Profiles

Normalization of gene expression profiles of a gene, or sets of genes, discussed here as mean SEPs (see RESULTS), implies that the mean over time of the profile equals zero, and the standard deviation equals one. This transformation allows direct and unbiased comparisons of expression profiles in the D and W groups of accessions, with a statistical evaluation performed at each time point (Figure 2 and Supplemental S-2 to S-4).

It is important to note that the differences in mean SEPs between D and W are produced by a set of genes that were likely affected by the domestication process (Figure 2). However, the large majority (21,666/22,427; 96.6%) of expressed genes did not exhibit significant differences in the mean expression profile (Supplemental S-4), implying that a large part of the transcriptome during fruit development was not affected by domestication.

In chili pepper, as in tomato, the first ten days after anthesis (DAA) are characterized by a period of very active cell division when the number of cell layers across the pericarp double, compared with that at anthesis, and this number is maintained through the end of development (Azzi et al., 2015). Fruit growth then proceeds into the cell expansion phase and continues until ∼40 DAA, progressing to full ripening at ∼50 DAA before entering senescence at ∼60 DAA (Martínez-López et al., 2014). During the period of cell expansion, from ∼10∼40 DAA, the dominant process is cell endoreduplication (Bourdon et al., 2010; Chevalier et al., 2011; Azzi et al., 2015). In *Capsicum*, pericarp thickness has a high positive correlation with the degree of polysomaty (Ogawa et al., 2010), which implies that accessions that have a thick pericarp have higher endoreduplication compared to accessions having a thin pericarp. This relationship has been corroborated through the induction of different ploidy levels (Ogawa et al., 2012).

The time lag observed for the peak of mean standardized expression between D and W accessions across the entire set of expressed genes (Figure 2) implies that domestication caused a shift in time and intensity of gene expression, favoring an earlier and higher expression maximum in D. Considering 463 genes that had the largest differences in expression profiles between D and W confirmed that the peak of expression for D occurs at 10 DAA compared to 20 DAA for W, and the peak expression value is significantly higher for D than for W. A similar expression pattern is seen for a gene encoding a G2/mitotic-specific cyclin, which is essential for control of the cell cycle at the G2/mitosis transition (Supplemental S-4). The transcript of this gene accumulated steadily during G2 and abruptly declined at mitosis. In *Arabidopsis*, which exhibits a similar trend in expression, members of the cyclin family are thought to be part of a developmental mechanism that coordinates the switch between proliferation and endoreduplication (Vanneste et al., 2011).

Here we found the peak gene expression for D at 10 DAA, a time when cell division is very active (Azzi et al., 2015). Furthermore, domesticated accessions bear substantially larger fruits than wild accessions (Paran and Van Der Knaap, 2007), and this larger fruit size is primarily achieved by increases in cell numbers (Guo and Simmons, 2011). As such, we grouped a set of 1,125 genes associated with cell reproduction by including genes annotated in 9 bioprocesses and examined the gene expression profiles (Supplemental S-5). Genes annotated for the cell cycle presented with an earlier increase and higher mean expression in D relative to W (Supplemental S-5). Given that both cell number and cell size, which are respectively determined by cell division and cell expansion (Gonzalez et al., 2007), contribute to fruit size, it is compelling that in the D accessions the cell division expression profile peaked earlier and higher than the W genotypes. This finding is consistent with that seen for tomato, in which cell division genes strongly influence fruit yield (Ariizumi et al., 2013).

The mean expression of the genes related to the cell cycle presented with a profile characterized by a large peak expression at 10 DAA for D, whereas W accessions had a smaller peak that occurred later at 30 DAA. We isolated a set of 542 genes that presented this pattern, which we termed “D10W30”. GO enrichment analyses produced a set of 86 biological processes (BPs) that were highly enriched among the D10W30 genes (Supplemental S-5 and SG). Besides 4 cell reproduction BPs, this set included 43 BPs associated with regulation of different cell processes including negative regulation of cellular process, and protein modification and transferase activity. Interestingly, such negative regulation has been associated with fruit development and ripening (Giovannoni, 2004). Other selected BPs include 4 that are related to cellular component organization, which has been linked to accumulation of soluble sugars and organic acids in fruits (Ma et al., 2019) and 3 that are related to cellular component assembly and also showed differential expression in a proteomic study of *Capsicum* (Guo et al., 2017). Another 3 in this set were identified with organelle fission or organization and 2 were related to microtubule-based process or movement, which are clearly associated with mitosis and have been linked to floral development in the genus *Aquilegia* (Voelckel et al., 2010) and autophagy BP that allows remodeling of intracellular structures during cell differentiation (Mizushima and Komatsu, 2011). In *Arabidopsis* this BP has been linked with the complete proteolysis of stromal proteins (Lee et al., 2013); see Supplemental S-5, S-6 and SG for details.

We plotted the expression profiles of genes that follow the D10W30 pattern, i.e., expression peaks on 10 DAA and 30 DAA for D and W accessions, respectively (Figure 2). Interesting examples this expression pattern are genes encoding: (i) the high mobility group B protein 6 (HMG B, 6), which belongs to a group of chromosomal proteins that regulate DNA-dependent processes and display a highly dynamic nuclear localization (Launholt et al., 2006); transcription factor ‘MYB-related protein 3R-1’, that in *Arabidopsis* synergistically maintain G2/M-specific genes repressed in post-mitotic cells and restricts the time window of mitotic gene expression in proliferating cells that has a role in determining organ size (Kobayashi et al., 2015); and (iii) the kinetochore protein NDC80, which is an essential component of the kinetochore complex that mediates chromosome segregation and spindle checkpoint activity to ensure proper cell division. In *Arabidopsis* the NDC80 mutant *mun-1* has a reduced cell division rate, aneuploidy and defects in chromosome segregation (Shin et al., 2018). See Figure 3 and Supplemental S-4.2.

The D10W30 set includes genes coding for “microtubule-associated protein TORTIFOLIA1”, and the “protein TPX2”, which have genomic fingerprints for domestication in *Capsicum* (Qin et al., 2014). TORTIFOLIA1 (TOR1) is a plant-specific microtubule-associated protein (MAP) that regulates cortical microtubule orientation and the direction of organ growth (Yao et al., 2008). TOR1 also determines microtubule organization by modulating microtubule severing (Wightman et al., 2013) and participates in organ elongation (Buschmann et al., 2004). On the other hand, TPX2 performs multiple roles in microtubule organization (Petrovská et al., 2013), such as regulating prospindle assembly before nuclear envelope breakdown (Vos et al., 2008) and is linked to fruit development in European pear (Nashima et al., 2013). Of the 300 domestication genes reported by Qin et al. (2014) and expressed during fruit development, 59 (∼20%) also showed significant (*P* < 0.05) differences between D and W accessions in this study.

An example of the coordinated time lag existent between D and W accessions is presented in Figures 3 and 4. The 14 genes involved in this network are pivotal for the cell cycle process, and thus functionally related, but also present a highly coordinated gene expression profile within D and W accessions, which markedly differs between these two groups by having the D10W30 expression pattern (Supplemental S-7, S-8 and S-9). It is important to assess the robustness of the links inferred between the genes in this network (Figure 3). With this aim we must take into account the fact that such links were inferred from ten fully independent datasets, i.e., the ones corresponding to each one of the 10 accessions. Then, if we assume that the selection of a gene to be part of the network has an error probability “*e*”, then the probability of selecting erroneously a gene to be part of the networks repeatedly in the ten accessions is *e*^-10^. Thus, even if the individual error probability is large, for example *e* = 0.1, the probability of having committed such error repeatedly in the ten accessions is vanishingly small; for *e* = 0.1 this equals 10^−10^, or one in ten billions.

Of the genes involved in the network of Figure 3A, two of them, (1) and (4) in the figure legend, encode two versions of the microtubule-associated protein 3. The *Arabidopsis* orthologous of these genes, *PLEIADE/AtMAP65-3*, has been shown to have physical and genetic interactions with the Transport Protein Particle II (TRAPPII), required to coordinate cytokinesis with the nuclear division cycle (Steiner et.al., 2016). Gene (2) in Figure 3A, labeled as DUF566 in the legend, contains the InterPro domain IPR007573 and corresponds with the *Arabidopsis* orthologous AT2G44190 (Table 2), which codes for the *endosperm-defective1* (*ede1*) gene; that encodes a microtubule-associated protein essential for microtubule function during the mitotic and cytokinetic stages that generate the endosperm and embryo, and thus is essential for seed formation (Pignocchi et al., 2009). Gene (3) in Figure 3, encodes a kinesin 3 which *Arabidopsis* ortholog, AT4G21270, encodes the *ATK1* gene, that has been demonstrated to be required for spindle morphogenesis (Chen et al., 2002). Consistently, the large majority of chili pepper kinesins follow the D10W30 expression pattern (Supplemental S-4.1). Gene labeled as “(5) − *POLLENLESS 3*” in the legend of Figure 3A, contains a tetratricopeptide repeat (TPR), and its closest *Arabidopsis* ortholog, AT4G20900, encodes the *TDM1* gene, which has been previously shown to be essential for meiotic termination (Cifuentes et al., 2016). Structural gene (6) in Figure 3A encodes shugoshin, a conserved kinetochore protein that prevents dissociation of cohesin from centromeres during mitosis (McGuinness et al., 2005). The closest *Arabidopsis* ortholog locus of this chili pepper sequence, AT3G44960, encodes 5 splicing variants of shugoshin, one of which has been reported to protect centromeric cohesion during meiosis (Kitajima et al., 2004); see also Supplemental S-9. It is intriguing that three of the chili pepper genes, the ones labeled as numbers (3), (5) and (6) in the legend of Figure 3A, have as closest orthologs in *Arabidopsis* genes reported primarily in meiosis (Chen et al., 2002), (Cifuentes et al., 2016) and (Kitajima et al., 2004), respectively, while the expression patterns of these *Capsicum* genes clearly correspond to genes involved in mitosis. This fact could be due to the large functional divergence between *Capsicum* and *Arabidopsis* genomes, which diverged from a common ancestor more than 150 million years before present (Qin et al., 2014).

On the other hand, the 8 TF candidates to be regulating the expression of the three structural genes in the network of Figure 3A (blue circles in Figure 3A; structural genes labeled as (1), (3) and (4) in the figure legend), those where assigned to the corresponding structural genes by a method that has been proved to successfully recover TF regulating the *AT3* gene in *Capsicum* (Arce-Rodríguez and Ochoa-Alejo, 2017; Zhu, et al., 2019; Sun et al., 2019); see METHODS and Supplemental S-8.

In summary, the functional network presented in Figures 3 and 4, demonstrates that domestication has produced a time lag in the expression of core cell cycle genes, anticipating for approximately 20 days the maximum expression of those genes and producing a higher standardized expression at the early fruit developing stage (10 DAA) in D when compared with W accessions.

Comparing gene expression profiles, rather than focusing on the differential expression of single genes at a given time, gives a better perspective on the complex interplay occurring in the transcriptome over time. Here, we demonstrated that a set of genes exhibits significant differences in expression profiles between D and W accessions during the development of chili pepper fruit. Genes in this set are associated with processes that involve cell regulation, cycle, localization, motility and assembly, as well as with autophagy and organelle organization. In particular, differences in time and intensity of gene expression of genes are related to cell reproduction, and provide an explanation at a molecular level of differences in fruit size between D and W accessions, which is the main morphological difference between these two genotype groups (Pickersgill, 2007).

## METHODS

### Statistical Design

RNA-Seq was performed as a factorial experiment with time (seven levels, 0, 10, 20, 30, 40, 50 and 60 DAA) and accession (10 accessions, 6 domesticated and 4 wild; see Table 1) as factors. The RNA-Seq library was the experimental unit, and two replicates of every combination of time per accession were independently replicated two times, for a total of 7 × 10 x 2 = 140 RNA-Seq libraries. After quality control, the raw reads were mapped to the *Capsicum* genome (CM334 v1.6) to obtain reliable counts for 22,427 genes. The relative expression of these genes is considered here as the output variable (see below).

### Plant Materials and Cultivation

Seeds of 10 *Capsicum annuum* accessions (Table 1) were surface sterilized with a 70% ethanol solution for 10 s before treatment with a 10% hypochlorite solution for 10 s and six rinses with distilled water. Wild accession seeds were similarly treated, after an initial treatment with 50% sulfuric acid solution to break seed dormancy. All accession seeds were germinated in plastic trays containing a mixture of three parts peat moss, one part perlite, one part vermiculite, one part sludge and two parts forest soil in a growth chamber, with 16 h light (photon flux of 70 μmol m^-2^ s^-1^) at 28 °C and 66% relative humidity. Three-week-old chili pepper plants were transplanted individually into plastic 5 L pots containing the same soil mixture described above. During transplantation, 15 g of a mycorrhizae fungal and beneficial bacteria mixture were added to optimize root growth and development. Plants were fertilized with Long Ashton solution every two weeks. Flowers and fruits at 0, 10, 20, 30, 40, 50 and 60 DAA were collected and immediately frozen in liquid nitrogen, and stored at −80 °C.

### RNA-Seq Library Construction and Processing

Total RNA was extracted from flowers and whole chili pepper fruits at different developmental stages using a NucleoSpin(tm) RNA Plant kit (MACHEREY-NAGEL) according to the manufacturer’s instructions. RNA was extracted from two biological samples comprising either flowers or fruits from 2-6 different plants. RNA quality was verified by determining the RNA Integrity Number (RIN) for each sample (Supplemental S-1).

Samples of total RNA were shipped to Novogene (https://en.novogene.com/) for library construction, sequencing and mapping to a reference genome. At Novogene, libraries were prepared and sequenced using the Illumina NovaSeq platform to obtain at least 20 million raw paired-end reads of 150 bp per sample. These reads were subjected to quality control and then mapped to the *Capsicum* reference genome CM334 v1.6 <u>(http://peppergenome.snu.ac.kr/).</u> Novogene provided the matrix of raw counts per library for each of the 35,883 *Capsicum* genes. These genes were identified by a protein product (when known), and annotated with Gene Ontology (GO) and Kyoto Encyclopedia of Genes and Genomes (KEGG) terms. In total, 140 libraries were processed (7 times x 10 accessions x 2 replicates of each combination of time x accession), yielding 2,298.8 million reads that mapped to the genome. The number of reads mapped to the genome had a minimum of 10.3, a median of 16.3, a mean of 16.4 and a maximum of 23.9 millions of reads per library. See Supplemental S-1 and SG.

### Estimation and Analyses of Standardized Expression Profiles (SEPs)

To avoid inclusion of genes that had very low or inconsistent expression patterns, only those genes having a raw count >0 in at least two of the replicates per accession were selected for analysis. After this filtering, 22,427 genes remained for analyses, and of these ∼62.5% were identified in the *Capsicum* genome. See Supplemental SG.

All results were maintained in an in-site MySQL (https://www.mysql.com/) relational database, and were analyzed with R version 3.4.4 (R Core Team, 2013). For each of the 10 accessions, we used the R package edgeR version 3.20.9 (Robinson et al., 2010) to obtain *P* values for each gene in the 6 contrasts between neighbor time intervals, i.e., 0-10, 10-20, 20-30, 30-40, 40-50 and 50-60 DAA. Following the method presented in Martínez-López et al. (2014), the gene expression tendency at each interval was classified as decreasing, steady or increasing, with adjustment of the Type I error to 0.01. For each gene within each accession, the mean standardized expression was calculated and the resulting 7-dimensional vector constituted the Standardized Expression Profile (SEP) in downstream analyses. There were 10 SEPs for each of the 22,427 genes consistently expressed during fruit development, with one for each of the accessions, and these SEPs were classified into the groups of interest: domesticated (D) with 6 elements, and wild (W) with 4 elements. To evaluate the difference between D and W SEPs, we calculated Euclidean distances between and within the groups and tested the hypothesis of equality of distances between and within SEPs with a t-test. Using a FDR (Benjamini and Hochberg, 1995) of 1% (0.01 in Q value) genes were classified as having an equal or distinct SEP in the D and W groups. For individual genes or groups of genes, we calculated the 95% CI for the means at each of the 7 time points sampled as well as the *P* value for the t-test of mean equality. A set of R functions was programmed to data-mine the results, and a binary file containing all data and functions is available upon request. Statistical procedures are described in detail in Supplemental S-2 to S-4.

### Network Estimation

We begin the network estimation with the 25 genes with expression profile D10W30 annotated with the cell cycle biological process in the 10 accessions. For all the 25⨯ (25-1)/2 = 300 different pairs formed by these 25 genes, we calculated Euclidean distance between their mean SEPs, selecting the pairs with a distance less than 1 standardized units. This stringent criterion selected only 10 gene pairs (3.3% of the total, and shown as edges between orange circles in Figure 3A), which include the 6 structural genes (orange circles in Figure 3A). The SEPs of these 10 gene pairs linked with black double headed arrows have a large (*r* > 0.96) and highly significant (*P* < 0.0001) mean Pearson’s correlation within the D or W genotypes. In contrast, the corresponding mean correlation between D and W accessions was small (*r* < 0.4) and not significant (*P* > 0.5), demonstrating that while these genes have a highly concordant expression within the D and W groups, the expression profiles are very different between those groups (Figure 3B and Supplemental S-7). Finally, the selection of the 8 TF candidates to be regulating the expression of the structural genes in the network of Figure 3A was performed employing an algorithm which select TFs with a highly concordant expression profile between TFs and each target gene (see Supplemental S-8 for details).

## ACCESSION NUMBERS

In process; All sequencing data will be delivered to the NCBI GEO data repository (https://www.ncbi.nlm.nih.gov/geo/).

## SUPPLEMENTAL INFORMATION

Supplemental (Supplemental.pdf) - Sections of this document are referred to in the text as ‘S-#’ where ‘#’ is the number of the corresponding section.

SG (SG.xlsx) - Supplemental Excel file with four sheets including information about genes and analyses of bioprocesses (BPs).

## FUNDING

This research was funded by the Consejo Nacional de Ciencia y Tecnología, México (Conacyt project 1570).

## AUTHOR CONTRIBUTIONS

Conceptualization, O.M and N.O-A; Methodology O.M, N.O-A and C.E-S; Formal Analysis, O.M, M.H.R-V and C.E-S; Investigation, M.L.A-R, F.H-G, F.C-H, C.H-K, F.G.R-M and C.E-S; Resources C.H-K, A.G-C and M.H.R-V; Data Curation O.M, C.E-S, M.L.A-R and F.C-H; Writing – Original Draft O.M, N.O-A and M.L.A-R; Writing – Review & Editing, O.M, N.O-A, J.J.O-O and C.E-S; Visualization O.M and C.E-S; Supervision O.M, N.O-A and M.L.A-R; Project Administration N.O-A; Funding Acquisition N.O-A, O.M, C.H-K, J.J.O-O, M.H.R-V and A.G-C.

## ACKNOWLEDGMENTS

F.C-H, C.E-S and F.G.R-M acknowledge Conacyt scholarships for PhD studies (numbers 707294, 630487 and 261122 respectively). We thank Dr. Victor Olalde-Portugal and M. Sc. Rosalinda Serrato-Flores in providing chili pepper seed germination and plant growth materials. The authors would like to thank Virgilio Alegría-Germán, Antonio Contreras, Dolores Ramírez, Juan B. Teran-Fraijo and María dS Fraijo-Encinas for helping us to collect the wild chili in Sonora. We also acknowledge Clara Borja-Castillo, Pamela Morín-Pérez, Jesús Gámez-Fernández, Alejandra Gómez-Elizarráz, José M. Villasuso-Aguiñaga, Elías López-Olvera, Sofía Martínez-Martínez, Isaac Aguirre-Manríquez, Jesús Caudillo-Corona, Frida Figueroa-Gómez and Valeria Gallardo-Onesto for help in sampling collection and processing. No conflict of interest declared.

## SUPPLEMENTAL

**NOTE**: Sections of this document are cited in the main text of the paper as “S-#”, where ‘#’ corresponds to the section in the table of Contents (below). In an effort to follow the standards of reproducible research (Peng, 2011), all relevant information is stored into a MySQL relational database named ‘SALSA’, and data as well as functions used in the analyses are in an R (R Core Team, 2013) binary object. These files are available upon request.

### S-1. Library sequencing and mapping to reference genome

As mentioned in the main text, after extraction we shipped the 170 total RNA samples to Novogene for quality control, sequencing and mapping to reference genome CM334 v1.6. The 170 samples correspond to 10 accessions (Table 1 in main text) *×*7 stages of fruit development (0, 10, 20, 30, 40, 50 and 60 DAA) *×* 2 independent biological replicates. Here we briefly describe and exemplify the procedures carried out in Novogene.

RNA sequencing was carried out in the Illumina NovaSeq platform, based on mechanism of SBS (sequencing by synthesis), and the sequencing workflow of the project is illustrated in Figure 1a, while Figure 1b shows the pipeline of the analyses and Figure 1c presents the quality control pipeline for the filtering of raw reads.

Original image data file from the Illumina sequencing platform were transformed into sequenced reads (raw reads) by CASAVA base recognition (Base Calling). Raw data are stored in FASTQ (fq) format files, which contain sequences of reads and corresponding base quality. In Figure 1c we see the post-processing of raw reads which consisted in (1) Remove reads with adaptor contamination, (2) Remove reads when uncertain nucleotides constitute more than 10 percent of either read (N *>* 10%), (3) Remove reads when low quality nucleotides (Base Quality less than 20) constitute more than 50 percent of the read.

Figure 2 presents examples of the results obtained from the library for sample ‘AS00R1’ (Replicate 1 of the time 0 DAA from accession AS); for brevity not all results are shown for this library and there are results for a total of 170 libraries, all of which were visually inspected before further processing. Figure 2a presents the plot of percentage of error rate (*Y*-axis) by position along the reads (*X*-axis), and in general, a single base error rate should be lower than 1%. Figure 2c shows the reads distribution to the reference genome as percentage of total raw reads, in categories (1) Adaptor related: (reads containing adapter) / (total raw reads), (2) Containing N: (reads with more than 10% N) / (total raw reads), (3) Low quality: (reads of low quality) / (total raw reads) and (4) Clean reads: (clean reads) / (total raw reads). For all 170 libraries the large majority of reads were in class (4), i.e., clean reads. Figures 2c and 2d refer to mapping the reads in the reference genome and will be commented below.

The algorithm for mapping filtered sequenced reads to the reference genome is shown in Figure 3.

In Figure 3 shows how the program HISAT2 was run with default parameters to map the clean reads to the genome. As examples of the result of the process Figure 2c shows the reads distribution to the reference genome by categories while Figure 2d shows the reads densities in chromosomes, in both cases for a single library, ‘AS00R1’ (Replicate 1 of the time 0 DAA from accession AS).

A total of 2298.8 millions of clean reads from the 170 RNA-Seq libraries were mapped to the genome, and the number of clean reads reads mapped to the genome per library ranges from a minimum of 10.3 millions up to a maximum of 23.9 millions with a mean of 16.4 millions.

To evaluate the accuracy of the results as well as the efficiency of the experimental procedures we can use the matrix of correlation coefficients between gene expression in samples. Figure 4 shows a partial view of that matrix for only 56 of the 170 libraries.

Correlation of the gene expression levels between samples plays an important role to verify reliability and sample selection, which can not only demonstrate the repeatability of the experiment but estimate the differential gene expression analysis as well. The closer the correlation coefficient is to 1, the higher similarity the samples are. Encode suggests that the square of the Pearson correlation coefficient, *r*, should be larger than 0.92 (under ideal experiment conditions). Correlation coefficients between samples indicates that the expression pattern is closer. In Figure 4 higher correlation coefficients, *r*, are represented by darker color, and replicates of libraries are set adjacent in both axis, while samples are ordered by accession at each axis. The higher correlation (*r* = 1; darkest color) is obviously present between each library with itself, which is shown in the main diagonal of the matrix. In Figure 4 samples are ordered at each axis by genotype (accession) and time (neighboring times are closer), and we can see a pattern of 4 *×*4 ‘squares’ corresponding to each one of the 4 accessions, the squares in the main diagonal correspond to correlations between each accession. In general data were highly consistent; in all cases correlations between replicates of the same accession and time were high and there was gradient from higher to lower correlations depending on time.

Novogene results also included all known Gene Ontology GO and Kyoto Encyclopedia of Genes and Genomes KEGG annotations of the *Capsicum* genome.

All results from Novogene were downloaded and kept in an in-site MySQL relational data base called ‘SALSA’.

**Figure 1.**
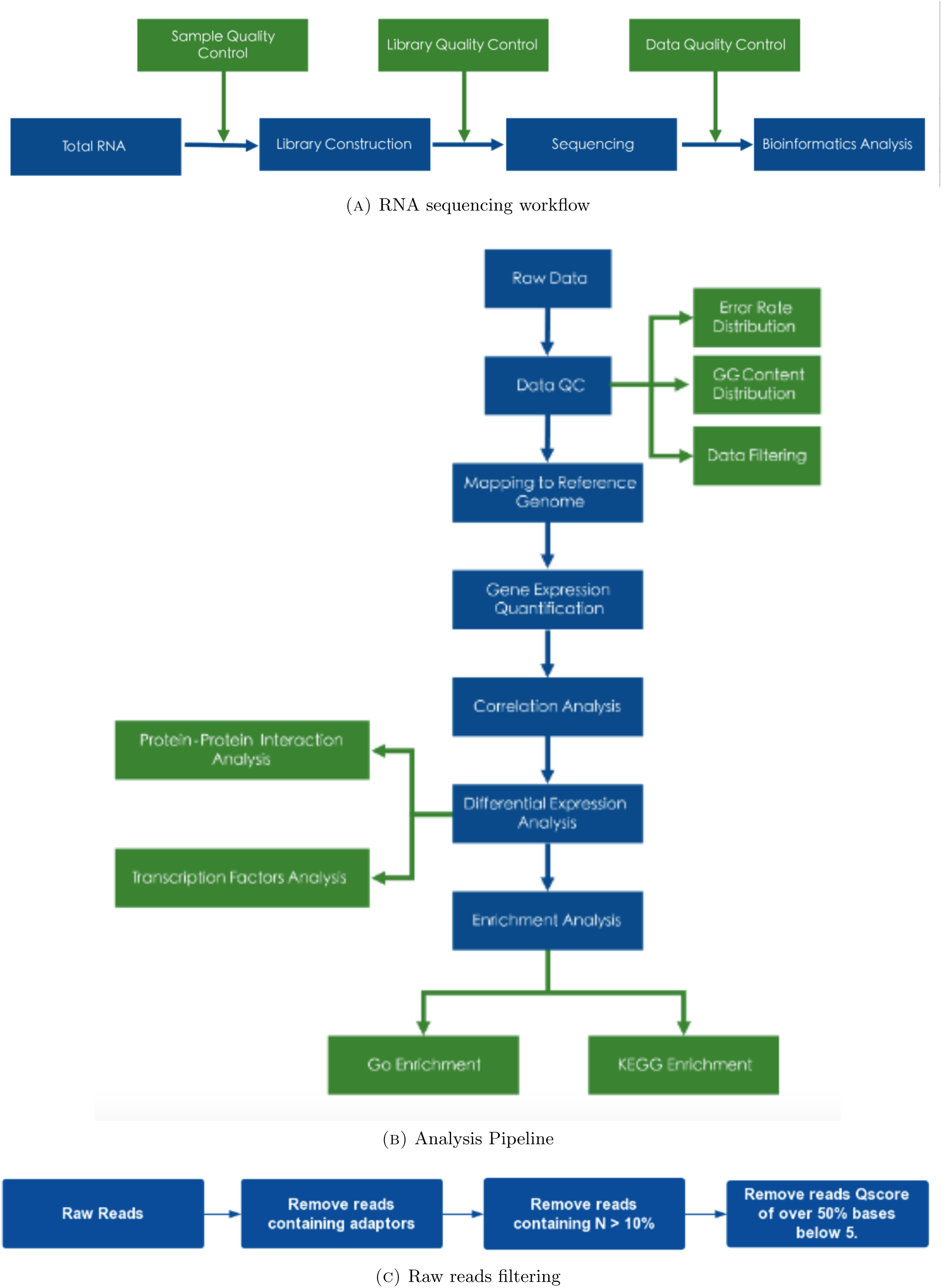
General procedure

**Figure 2.**
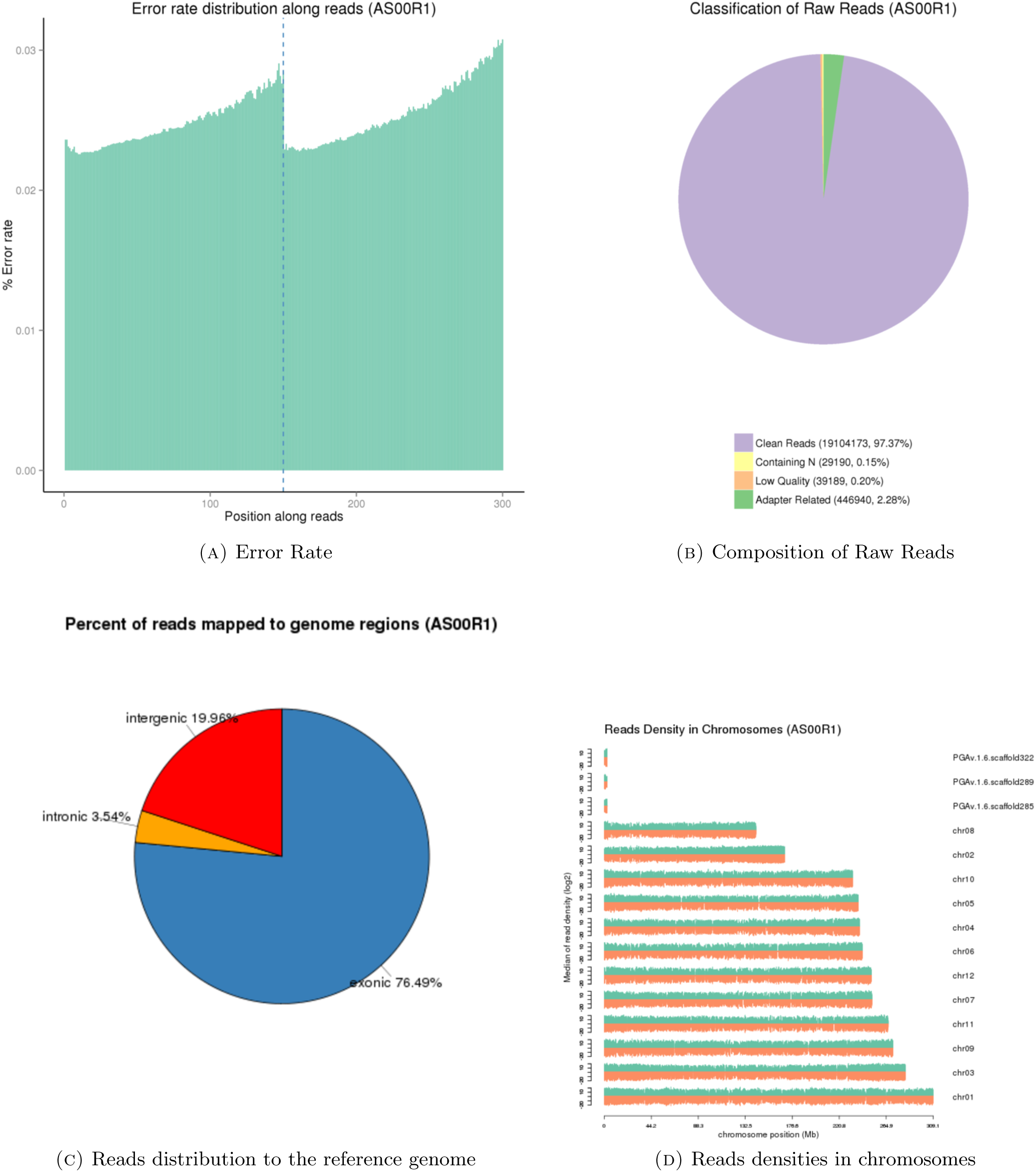
Examples for the library obtained from sample ‘AS00R1’ (Replicate 1 of the time 0 DAA from accession AS).

### S-2. Standardized Expression Profile (SEP) estimation

The majority of RNA-Seq studies (Wang et al., 2009) are focus on the direct estimation of differential gene expression. However, in our case we want to estimate the *expression profile*, i.e., the change of the relative gene expression through time. Given that our experiment was an RNA-Seq time-course (Luan and Li, 2003; Iglesias-Martinez et al., 2016) study, the emphasis was to summarize the changes that occur from one point in time to the next. We sampled seven times during fruit development, say *t*_1_, *t*_2_, *…, t*_7_, corresponding to 0, 10, 20, 30, 40, 50 and 60 DAA, thus the contrasts of interest were between times *t*_*i*_, *t*_*i*+1_; *i* = 1, 2, *…*, 6; i.e., between the six neighboring intervals. Let’s denote the true mean gene expression for a given gene within one of the accessions as *µ*_*i*_; *i* = 1, 2, *…*, 7. Then, for each neighbor interval we had three possibilities, say, gene expression decreases from time *i* to time *i* + 1, denoted as ‘D’ and expressed by the hypothesis *µ*_*i*_ *> µ*_*i*+1_; steady gene expression from time *i* to time *i* + 1, denoted by ‘S’ and corresponding to *µ*_*i*_ = *µ*_*i*+1_ and finally gene expression increases from time *i* to time *i* + 1, denoted as ‘I’ corresponding to *µ*_*i*_ *< µ*_*i*+1_. To decide between these alternatives we employed the program edgeR (Robinson et al., 2010) as described below.

**Figure 3.**
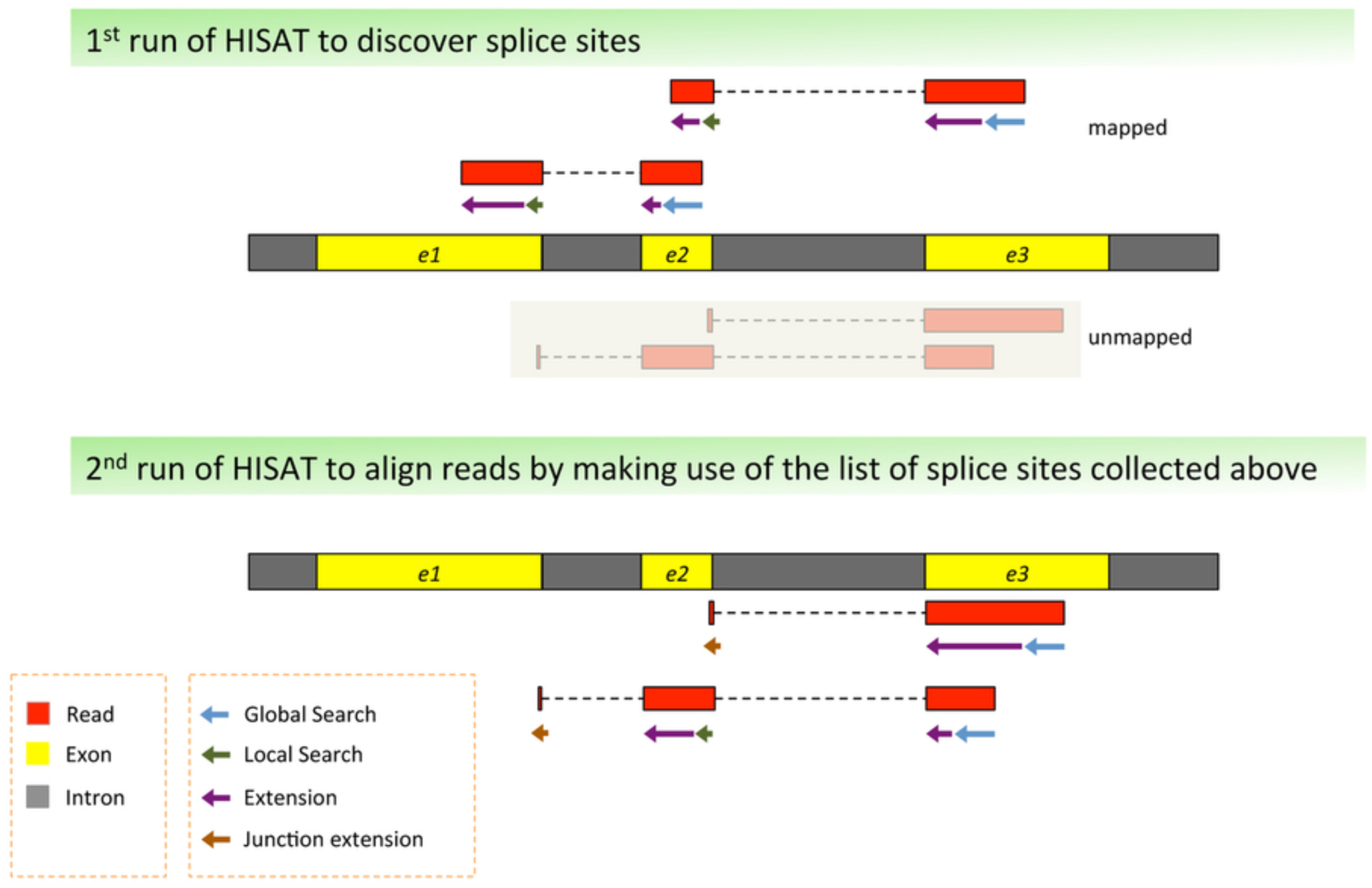
Mapping process

**Figure 4.**
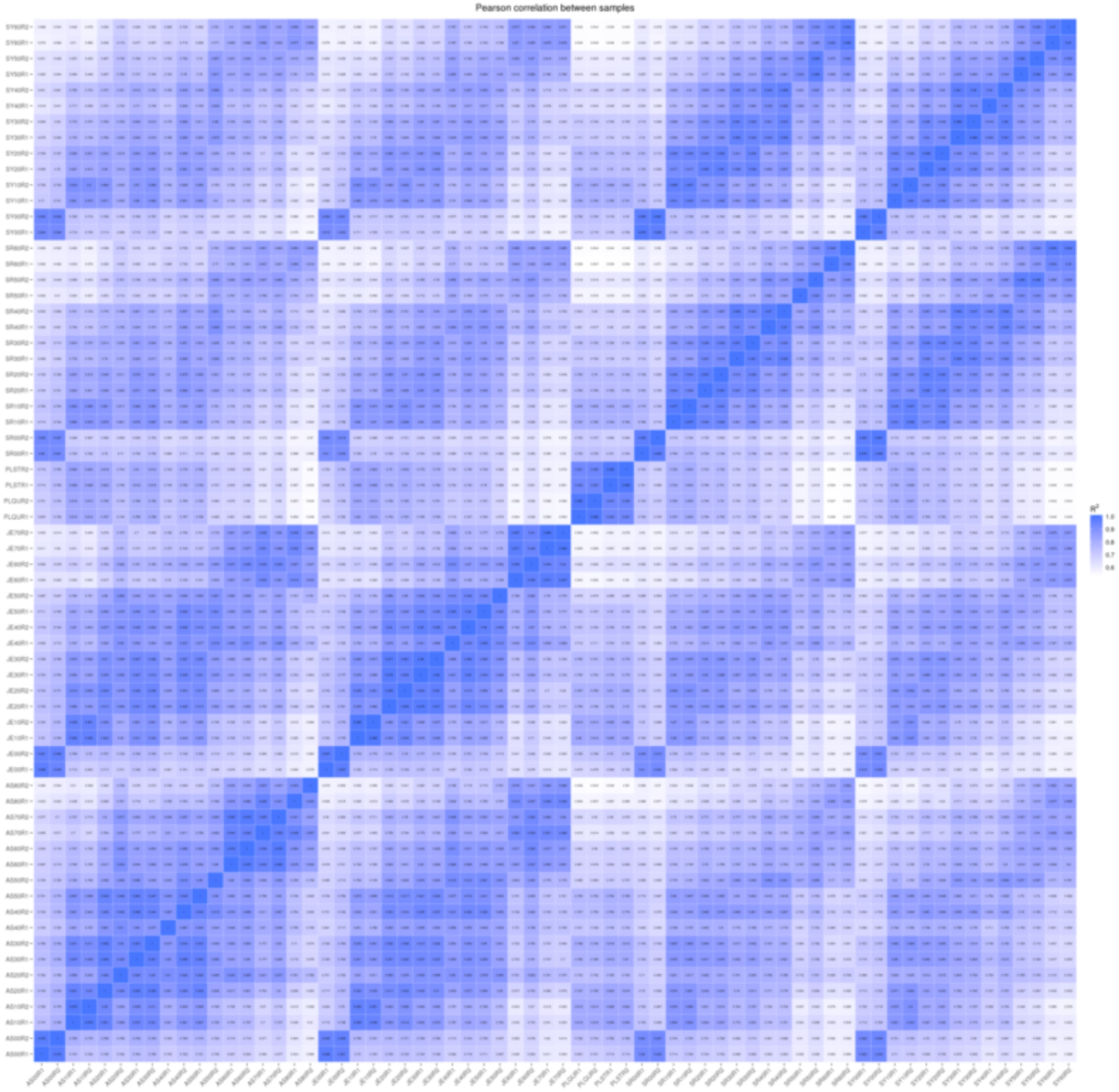
Matrix of correlation coefficients between samples (replicates are adjacent).

It is important to realize that we want to statistically summarize a gene expression profile that exist in the six dimensional space created by the contrasts at neighbor intervals, and thus six tests of hypothesis, one for each one of the neighboring intervals, need to be performed. Given that we test multiple hypotheses (one for each interval), we need to consider the Bonferroni correction (Abdi, 2007) for the probability of calling two expression profiles as statistically different. Thus, to obtain an approximate probability of Error Type I, *p*^*\**^, when performing 6 tests, we need to use a *p*^*\**^ value equal to *p*^*\**^ = *p*^6^, where *p* is the value employed at each one of the 6 individual tests. In our case we fixed *p\** to be equal to 0.01 or 1%. Note that we were not going to directly perform or use hypothesis tests between different gene expression profiles, but only use the expression profile as a reasonable summary of gene expression through time. The basic idea behind this method of estimation was previously published by our group in Martí nez-López et al. (2014).

To obtain the *p*^*\**^ values needed by the method, we run edgeR (Robinson et al., 2010) on the matrix of raw counts of reads for each one of the accessions, performing the tests for each gene in contrasts *t*_*i*_ *vs. t*_*i*+1_, *i* = 1, 2, *…*, 6, i.e., for the differences in expression between the 6 pairs of neighboring intervals.

Because at each time interval we had three possibilities for the change of gene expression, as said before, ‘D’ when *µ*_*i*_ *> µ*_*i*+1_; ‘S’ when *µ*_*i*_ = *µ*_*i*+1_ and ‘I’ when *µ*_*i*_ *< µ*_*i*+1_, we call the realization of these profiles ‘Ternary Models’, because only 3 possibilities were contemplated at each one of the neighboring intervals. Ternary Models can be represented by the six successive results obtained in the intervals; for example, model ‘SSSSSS’ represent the case where gene expression was steady, i.e., with no significant change during all fruit development, while model ‘DDISS’ denotes the case where expression decreased from 0 to 10 and 10 to 20 DAA, then increased from 20 to 30 DAA and then stayed steady in the last two intervals, from 40 to 50 and 50 to 60 DAA. Thus, by counting all possibilities we had a total of 3^6^ = 729 different Ternary Models.

To obtain raw estimated expression profiles we calculate, for each gene within each accession, the mean gene expression of the two biological replicates in FPKM units (Mortazavi et al., 2008)^1^. This gave a vector of seven numbers, say **m** = (*m*_1_, *m*_2_, *…, m*_7_), corresponding to the seven times points where the expression was estimated. The algorithm to obtain the Ternary Model profile from the raw estimated expression profile, **m**, needs also the 6-dimensional model vector

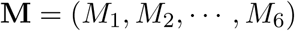

which contains the letters that denote the change at each one of the 6 intervals; i.e. *M*_*i*_ *∈ {D, S, I}*; *i* = 1, 2, *…*, 6, i.e., the Ternary Model for the gene.

The algorithm to calculate the Ternary Model profile is presented in the next list.

#### Algorithm to obtain the Ternary Model profile ‘o’ from input {**m, M**}

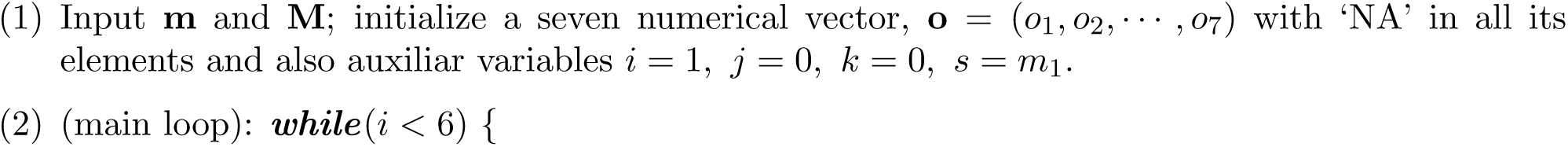
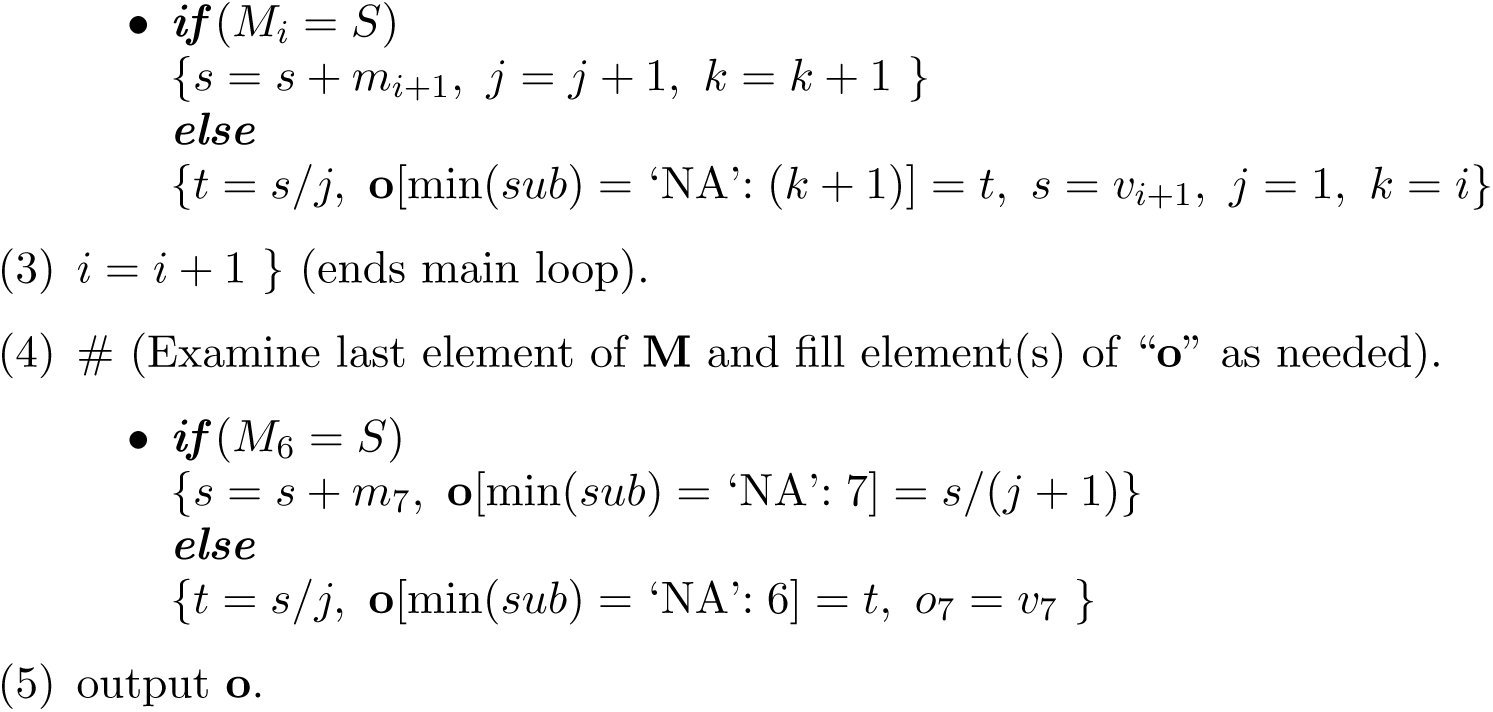

In the algorithm the elements of the output vector “**o**”, denoted by “**o**[min(*sub*) =‘NA’: *x*]”, are all elements of the vector that were ‘NA’ from the smallest subindex (*sub*) to *x*. The algorithm to calculate the Ternary Model profile obtains a vector, **o**, in which the values of steady intervals (intervals with ‘S’ in the model) are fill with the average of the corresponding values of the elements of **m**. This is so because when there was not statistical significant changes in one or more intervals, the best estimate of the mean expression is given by the average of the corresponding values of **m**.

A pair of numerical examples illustrate this algorithm, which converts a raw estimated expression profile, **m**, into the vector, **o**, which includes the Ternary Model information, **M**.

Firstly, consider the case of the gene with id=3 in accession AS; for this gene we have **M** = ‘SSSSSS’ (no interval with a significant change) and the rounded numerical values of the raw estimated expression profile are

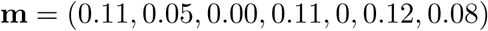

Because none of the changes in expression between neighboring intervals are significant (model is ‘SSSSSS’), all seven values of expression are averaged to obtain each one of the the seven values in **o**, say

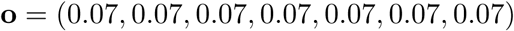

Secondly, consider a gene with a more interesting Ternary Model, say for example gene with id=526 in accession AS, which has **M** = ‘DSSISS’. This gene decreases from 0 to 10, stays steady from 10 to 30, increments from 30 to 40 and then remains steady up to 60 DAA. The rounded numerical values of the raw estimated expression profile are

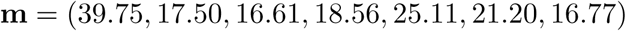

Applying the algorithm to this vector we obtain

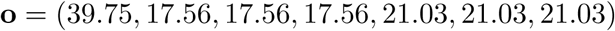

In this case we have *o*_1_ = *m*_1_ = 39.75 because in the first interval, *M*_1_ = ‘D’, we had a significant decrement from the expression at 0 DAA, 39.75, to the expression at 10 DAA, 17.50, but such decrement was followed by two steady states (**M** = ‘DSSISS’). Now, note that from 10 DAA up to 30 DAA expression was steady, i.e., *M*_2_ = *M*_3_ = ‘S’, thus the values of *o*_2_, *o*_3_ and *o*_4_ are obtained as the average of the values in *m*_2_, *m*_3_ and *m*_4_, i.e., the average of 17.50, 16.61 and 18.56 which equals 17.56, thus *o*_2_ = *o*_3_ = *o*_4_ = 17.56. In interval *M*_4_ (from 30 to 40 DAA) we have a significan increment, from *m*_4_ = 18.56 to *m*_5_ = 25.11, but such increment was followed by two steady intervals, *M*_5_ = *M*_6_ = ‘S’, and thus values of *o*_5_, *o*_6_ and *o*_7_ are equal to the average of 25.11, 21.20 and 16.77 which is 21.03.

Note that vectors of expression profiles, **o**, are not standardized to have a mean over time of 1 and a standard deviation of 1. Thus the last step to obtain Standardized Expression Profiles (SEPs) is to subtract the mean and divide by the standard deviation all elements of **o**, say to obtain the SEP, **s** from **o** we standardize setting *n*_*i*_ = (*o*_*i*_ *−* **ō**)*/S*_**o**_, where **ō** is the average of the seven elements of **o** and *S*_**o**_ is the standard deviation of the elements of **o**.

In the second example (gene with id=526 in accession AS) we have that **ō** = 22.21 and *S*_**o**_ = 7.93, thus the final representation of the Standardized Expression Profile (SEP), **s**, is given by

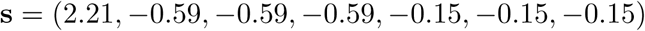

which has an average of 0 and a standard deviation of 1, i.e. it is ‘standardized’.

In summary, the estimation of a SEP, **s**, proceeds following the steps *{***M, m***}⇒* **o** *⇒* **s**, and it takes into consideration the mean gene expression **m**, which for each time is the average resulting from two RNA-Seq libraries as well as the statistical significance between neighboring times, contained in the Ternary Model **M**, adjusting the expression at each time to reflect significant changes by averaging the expression intervals where there is not significance, obtaining the Ternary Model profile, **o**, to finally obtain the SEP, **s**, by standardizing **o**. Even when this procedure could be judged as highly convoluted, it has a great advantage: It allows to compare gene expression profiles throughout time independently of the raw gene expression and it integrates the available statistical evidence for expression change between neighboring times.

Figure 5 shows the plot of the SEP for gene with id=526 in accession AS, presented before as second example.

**Figure 5.**
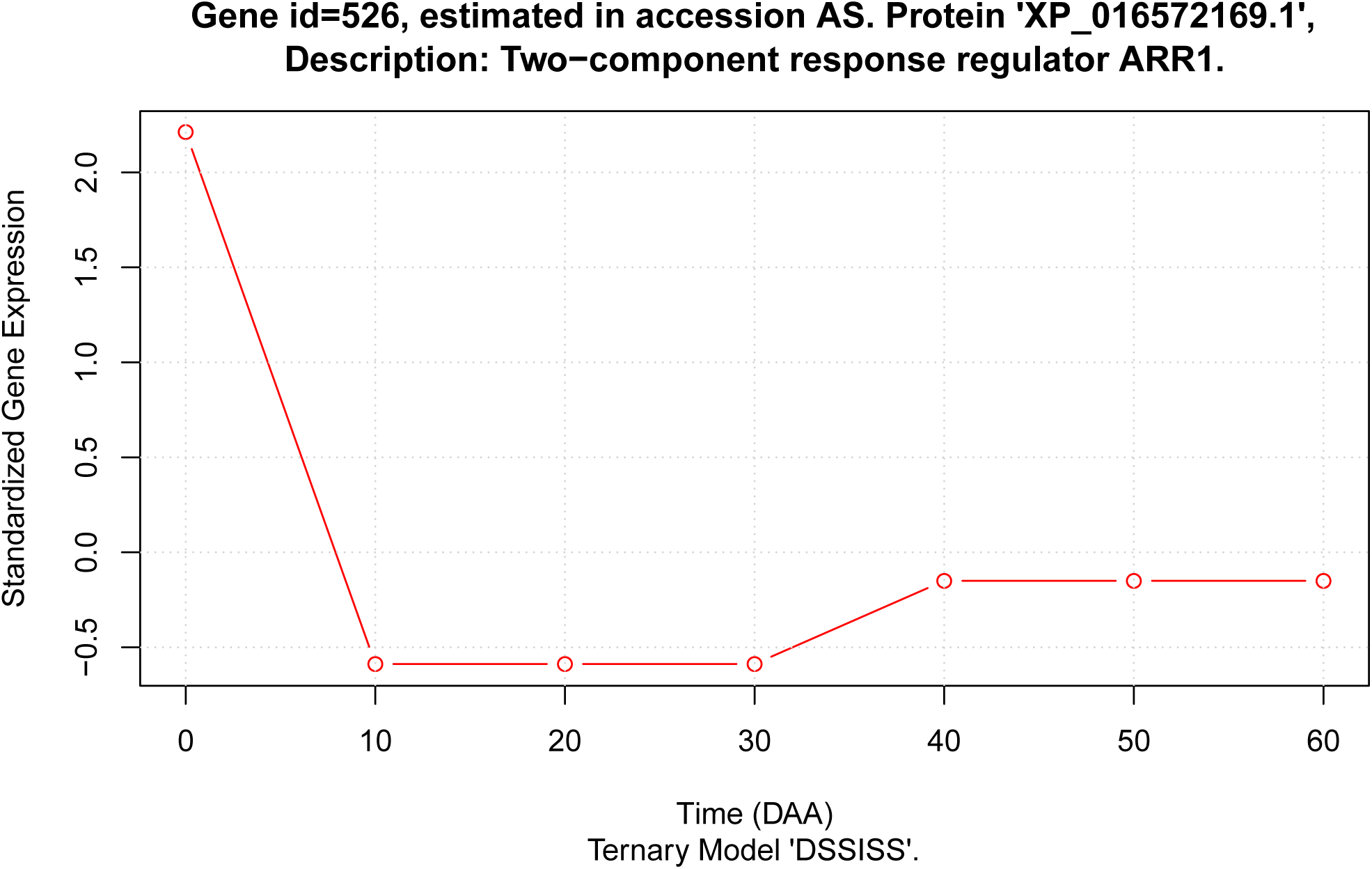
Example: Standardized Expression Profile (SEP; **s** vector) for gene with id=526 in accession AS.

In Figure 5 we can appreciate the plot of the SEP used as second example. Additionally to showing the best estimates of standardized changes in expression throughout time, we can see how the SEP model preserves the relative magnitude of expression changes; observing this plot we can immediately notice that the change from 0 to 10 DAA, with a total absolute difference of 2.8 standardized units, is much larger than the change from 30 to 40 DAA, which has a total absolute difference of 0.74 standardized units.

### S-3. Testing differences between Domesticated (D) and Wild (W) SEPs

The focus of this work was the detection of changes in standardized expression profiles (SEPs) between D and W accessions caused by the domestication process. We studied 10 accessions, 6 D and 4 W (see Table 1 in the main text), and found that a total of 22427, representing approximately 64% of the genes annotated in the *Capsicum* genome (CM334 v1.6) were consistently expressed in all 10 accessions at one or more of the times sampled and in more than one of the two biological replicates per accession.

For each gene we had 10 SEPs, 6 from D and 4 from W accessions, and we want to discriminate with a univariate statistic if there were differences in SEPs when grouping them in the D and W sets. For this we selected the Euclidean distance between SEPs, defined as

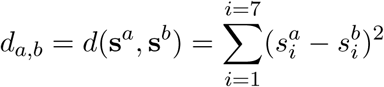

where **s**^*a*^, **s**^*b*^ are two different profiles for the same gene. For a given gene we calculated the total of 10(10−1)*/*2 = 45 different distances, *d*_*a,b*_; *a* ≠ *b*, and classified those distances into two groups, *distances between* D and W accessions and *distances within* one of the groups. The number of distances between D and W is equal to 6 *×* 4 = 24, while the remaining 45 *−* 24 = 21 distances happen within the two groups, say 6(6 *−* 1)*/*2 = 15 within D accessions and 4(4 *−* 1)*/*2 = 6 within W accessions.

For a single gene, our interest was to detect significant differences in SEPs between the D and W accessions, and this can be translated to the statistical hypothesis *ℋ*_0_ : *µ*_*b*_ = *µ*_*w*_ *versus H*_*a*_ : *µ*_*b*_ *> µ*_*w*_, where *µ*_*b*_ and *µ*_*w*_ are the true means of the distances between and within the D and W groups, respectively. If we accept the null hypothesis *ℋ*_0_ as true, then we have no evidence of differences between SEPs in the D and W accessions, while if this hypothesis is rejected in favor of *ℋ*_*a*_ : *µ*_*b*_ *> µ*_*w*_ (note that this alternative implies a one-tail test), we conclude that the mean distance between the two groups is significantly larger than the distance within those groups, and this implies a difference in SEPs between D and W. To perform the statistical test we assayed a randomization test comparing it with the usual parametric one tail t-test, and found that those alternatives were almost equivalent, opting for the second given the high computational cost of the second and the large number of tests (22427) that needed to be performed.

Figure 6 presents the histogram of the *P*-values obtained in the 22427 test of the null hypothesis *ℋ*_0_ : *µ*_*b*_ = *µ*_*w*_ *versus ℋ*_*a*_ : *µ*_*b*_ *> µ*_*w*_.

An interesting feature in Figure 6 is that the first bar, including *P* values between between 0 and 0.05, includes 4465 cases, approximately 20% of the total. This indicates that the *P* distribution of the tests performed is far from being uniform, as expected from randomized tests (Bland, 2013). And because we tested all genes expressed during fruit development, the non-uniformity of the *P* distribution for the tests implies that selection had an important role in the modification of SEPs.

Table 1 presents the matrix of average mean distances between 22427 SEPs, corresponding to equal number of genes, in the 10 accessions.

In Table 1 we can see that the minimum of the mean distances, 1.63 (in blue), occurs between SR an SY, two W accessions, while the maximum, 2.40 in red, happens between AS and SR as well as between AS and ST, in both cases a D and W accessions respectively. On the other hand, the mean average distance within D and W accessions (21 values from the matrix) is 2.02, while the mean average distance between D and W accessions (24 values from the matrix) is 2.18; i.e., the D and W accessions form two well segregated groups.

The dendrogram presented in the Figure 1 of the main text was obtained by applying the agglomerative Ward’s algorithm on the distance matrix shown in Table 1. In that figure W accessions are grouped in a single cluster (left hand side), well separated at a mean Euclidean distance *>* 2.8 from the one formed by the 6 D accessions (right hand side), this shows that gene expression variability within the W and D groups is smaller than the distance between those groups.

**Figure 6.**
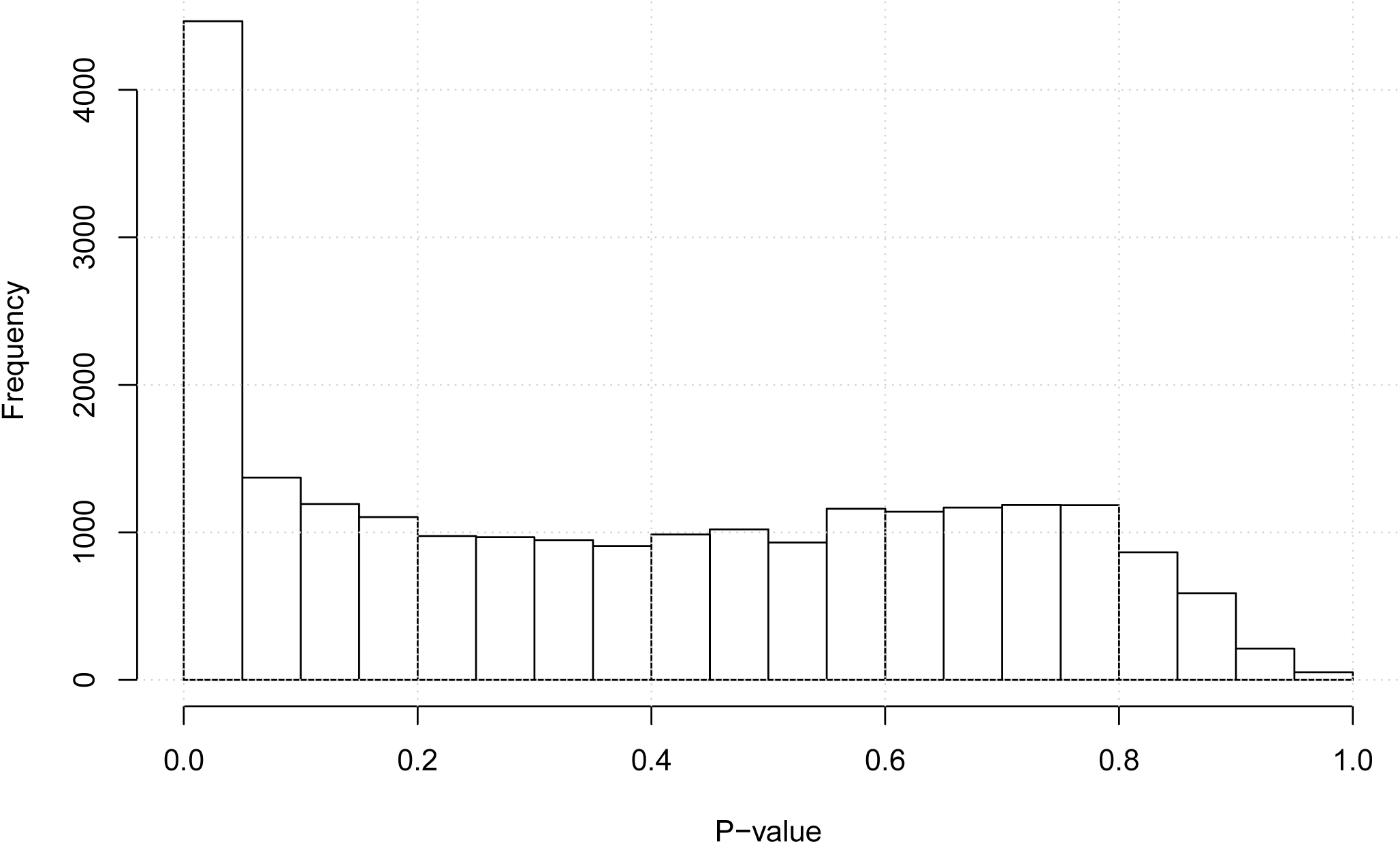
Histogram of the *P*-values obtained in the 22427 test of the null hypothesis *ℋ*_0_ : *µ*_*b*_ = *µ*_*w*_ *versus H*_*a*_ : *µ*_*b*_ *> µ*_*w*_ employing the one-tail t-test.

**Table 1.** Matrix of average mean distances between the SEPs in the 10 accessions.

|  | CM (D) | CO (W) | CW (D) | JE (D) | QU (W) | SR (W) | ST (D) | SY (W) | ZU (D) |
| --- | --- | --- | --- | --- | --- | --- | --- | --- | --- |
| AS (D) | 2.13 | 2.33 | 2.05 | 1.91 | 2.35 | 2.40 | 2.40 | 2.37 | 2.27 |
| CM (D) |  | 1.96 | 2.07 | 2.01 | 1.98 | 1.95 | 2.11 | 1.97 | 1.98 |
| CO (W) |  |  | 2.26 | 2.22 | 1.80 | 1.78 | 2.18 | 1.74 | 1.95 |
| CW (D) |  |  |  | 2.02 | 2.29 | 2.31 | 2.23 | 2.31 | 1.97 |
| JE (D) |  |  |  |  | 2.26 | 2.29 | 2.31 | 2.18 | 2.08 |
| QU (W) |  |  |  |  |  | 1.91 | 2.15 | 2.00 | 2.12 |
| SR (W) |  |  |  |  |  |  | 2.21 | 1.63 | 2.05 |
| ST (D) |  |  |  |  |  |  |  | 2.23 | 2.06 |
| SY (W) |  |  |  |  |  |  |  |  | 2.04 |

### S-4. Analyses per time of SEPs in D and W accessions

For each one of the 22427 genes expressed during fruit development we have 10 SEPs, and in the previous section we have described the univariate test performed on the Euclidean distances to decide if the SEPs in the set of 6 D accessions could be considered different to the 4 ones in the W group. Independently of the fact that SEPs grouped into the D and W could be considered to be equal or not by that test, we can additionally analyze the differences between SEPs in the 7 stages of development (0, 10, 20, *…*, 60 DAA), grouping a single gene or sets of genes in the D and W sets.

Let’s denote as 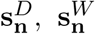, the 7-dimensional SEP vectors for genes in an arbitrary set of genes **n**, which cardinality is *n*, i.e., the set **n** is constituted by *n* different genes (|**n**| = *n*).

As an example, define **n** as the set formed with the gene with identifier 580 (a single gene). Then 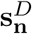 is constituted by 6 different vectors, each one corresponding to each one of the 6 D accessions, while 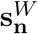 is formed by 4 different vectors, each one corresponding to each one of the 4 W accessions. Now, for each stage of development, *i* = 1, 2, *…*, 7, we have two sets of independent standardized gene expressions, say, *d*_*i*_ = *{s*_*ij*_*}*; *j* = 1, 2, 3, 4, 5, 6 for D and *w*_*i*_ = *{s*_*ik*_*}*; *k* = 1, 2, 3, 4 where the subindex *j* denote accession, D or W, respectively. Note that all elements *{s*_*ij*_*}, {s*_*ik*_*}* are fully independent, because each one of them was estimated from a different RNA-Seq library.

For each one of the stages of development, the hypotheses of interest are: 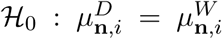 *versus*: 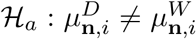, where *i* = 1, 2, *…*, 7 and : 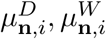 represent the true means of standardized expression at developing stages 0, 10,, 60 DAA, respectively. The number of standardized observations in the sets D and W depend on the number of genes in the set **n**, as before |**n**|= *n*, thus if *n* = 1 (a single gene tested), then the number of observations to be included in the two sets to be tested are 6 for D and 4 for W, while in general for any any set of genes **n** with *n* genes we will have 6*n* and 4*n* observations for D and W, respectively. To perform the tests 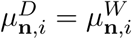, *i* = 1, 2, ...,7 as well as to obtain 95% Confidence Intervals (CIs) for the means of each group at each one of the times we employed the two tail t-test. The procedure to perform the test and plot the results for any arbitrary set of genes was programed in an R function.

On the other hand, it was considered important to evaluate the stage at which the maximum expression of a gene was reached. In this case for each SEP we determine stage (0, 10, *…*, 60) at which the maximum standardized expression is reach. Denote as *m*_*i*_ the point of development at which the maximum of the SEP vector **s**_*i*_ = (*s*_*i*1_, *s*_*i*2_, *…, s*_*i*7_) is found. For example, if max(**s**_*i*_) = *s*_*i*3_, this means that the maximum standardized expression took place at the third stage (*i* = 3), corresponding to 20 DAA, thus the value of *m*_*i*3_ is 20, etc. For any gene or set of genes **n**, we calculated the set of maxima in D and W accessions and tested the hypothesis 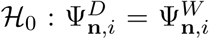 *versus* 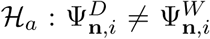, where 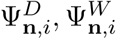 represent the true means of the maximum standardized expression and calculated the corresponding 95% CI.

The functions to analyze and plot the results for an arbitrary set of genes, **n**, where employed to obtain figures 2, 3 and 4 presented in the main text. In these, as in any results from such functions, the corresponding plots show the 95% CI for mean standardized expression as thin lines at each stage of development, while the estimated mean maximum expression is shown by asterisks with their corresponding 95% CIs shown by an horizontal line. To illustrate these kinds of results we present examples for two genes.

Our first example corresponds to the results obtained for the gene with id=580, and plots are presented in figures 7 and 8.

Figure 7 presents SEPs for the *Capsicum* fibrillin (FBN). Fibrillins are nuclear-encoded, plastid proteins associated with chromoplast fibrils and chloroplast plastoglobules (Singh and McNellis, 2011), and in Figure 7 we can appreciate how expression of FBN is highly concordant in all accessions. In that figure the points plotted are slightly displaced in the *X* axis (DAA) to avoid line and symbols overlapping. In all accessions TMs for FBN had a low standardized expression from 0 up to 40 DAA, where the expression increases rapidly, reaching the maxima at 50 (in 3 accessions; 2 D and 1 W) or 60 (7 accessions; 5 D, 3 W) DAA. The FBN gene does not present a significant difference in distances between D and W accessions, having a *P*-value of 0.8 in that test, and exemplifying a case of a gene which was not affected by domestication. On the other hand, Figure 7 presents mean SEPs for the FBN gene. That figure was produced with our function `‘TMmean.plot()’`, which also produced the output presented in Appendix S-11.

In Appendix S-11 we see that the results include tables of means and CI for the means for the standardized expression at each point in time; those CI are plot as thin vertical lines in Figure 8, allowing the visual judgment of the difference between the means in the D (red) and W (blue) sets. For all time points (0, 10, *…*, 60 DAA) we see that the CI of D and W overlap, and the lack of a significant difference can be observed in the *P*-values for the t-test of means D vs W per time point in Appendix S-11. The second analysis performed is the estimation of means and t-test for the maxima in the D and W groups. Appendix S-11 presents the means and 95% CI for those estimates. The mean for the D group is 56.67 DAA while the mean for the W set is only slightly different, 57.5, with CIs overlap between the two groups. Finally, the lack of significance of the difference in the mean maxima between the two groups is confirmed by the t-test, which gives a value of *P* = 0.8065 The function even gives the interpretation of the result in the line: ‘(Genes are Early in D but the difference is NOT significant at 0.05)’. Figure 8 presents the means of the times where the maximum expression for each set is estimated as asterisks and the corresponding CI as broad horizontal lines. From all the analyses we can conclude that the FBN gene has a highly similar expression pattern in both, D and W accessions. This kind of analysis and plots were used for figures 2, 3 and 4 in the main text with different sets of genes.

**Figure 7.**
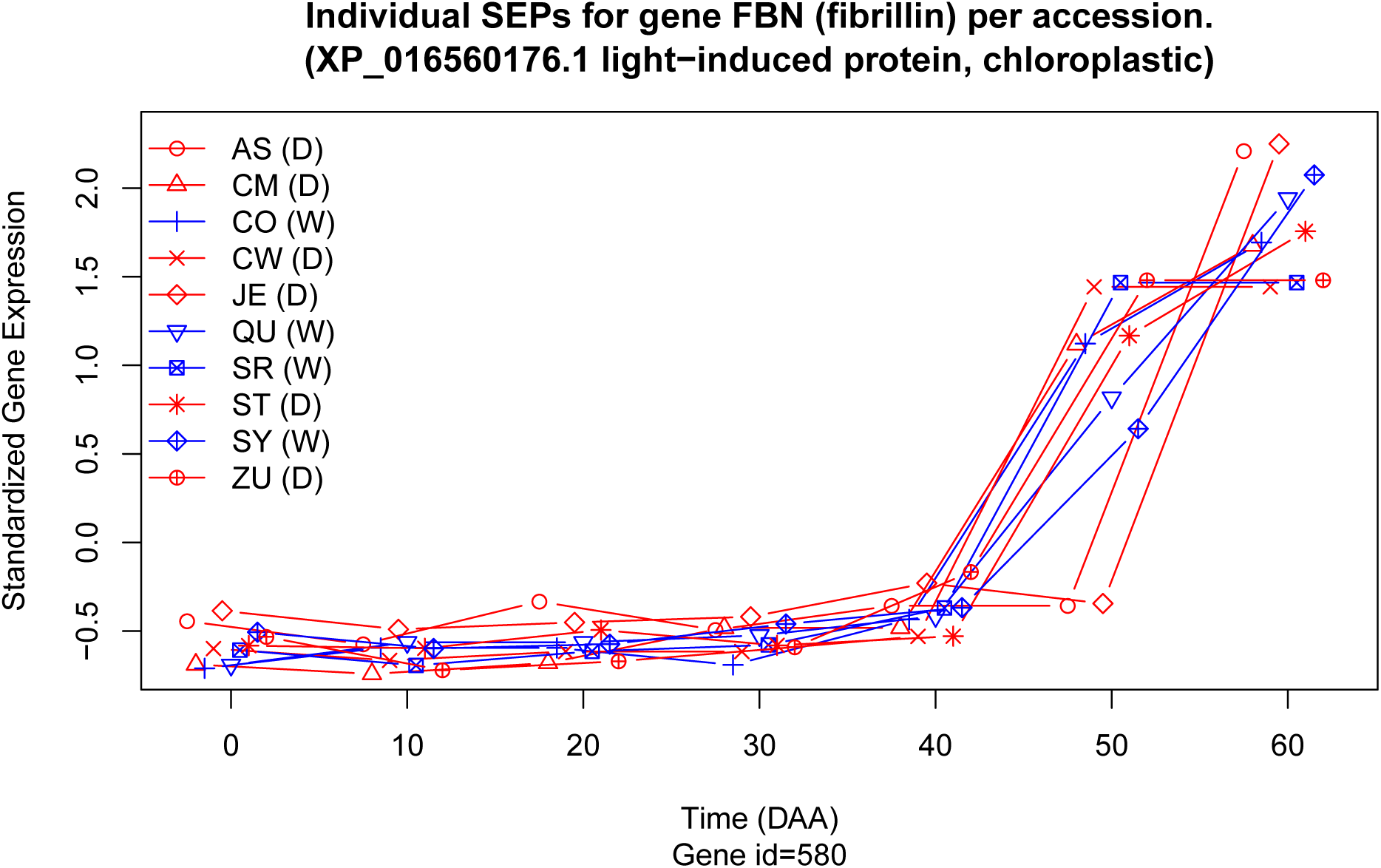
SEPs per accession for a gene with highly concordant expression patterns in all 10 accessions. Values per accession were slightly displaced in the *Y* axis to avoid overlapping.

Figures 9 and 10 present plots for a gene with highly different SEPs between D and W and Appendix S-11.1 presents the statistical analysis for this case.

The gene with id=19147, a transcription factor identified as ‘B3 domain-containing protein At5g42700-like’ and with protein identifier XP 016568750.1, was highly significant (*P <* 4.6*×* 10^*−*14^) in the univariate test for differences in SEPs between D and W, and in fact Figure 9 shows that this gene has SEPs which in D accessions have a maximum at 10 DAA, while in W the maximum is present at 30 DAA. This expression pattern indicates that this gene belongs to the group of ‘D10W30’ genes defined in the main text. Indeed, in Figure 10, which presents the mean SEPs for the gene and the 95% CIs for time of maximum expression over the the *X* axis, and standardized gene expression over the *Y* axis, shows that the maxima are different for D and W, while there are significant differences in mean expression at 10, 30, 40, 50 and 60 DAA. Appendix S-11.1 presents the R output with the statistical results obtained in the analyses.

**Figure 8.**
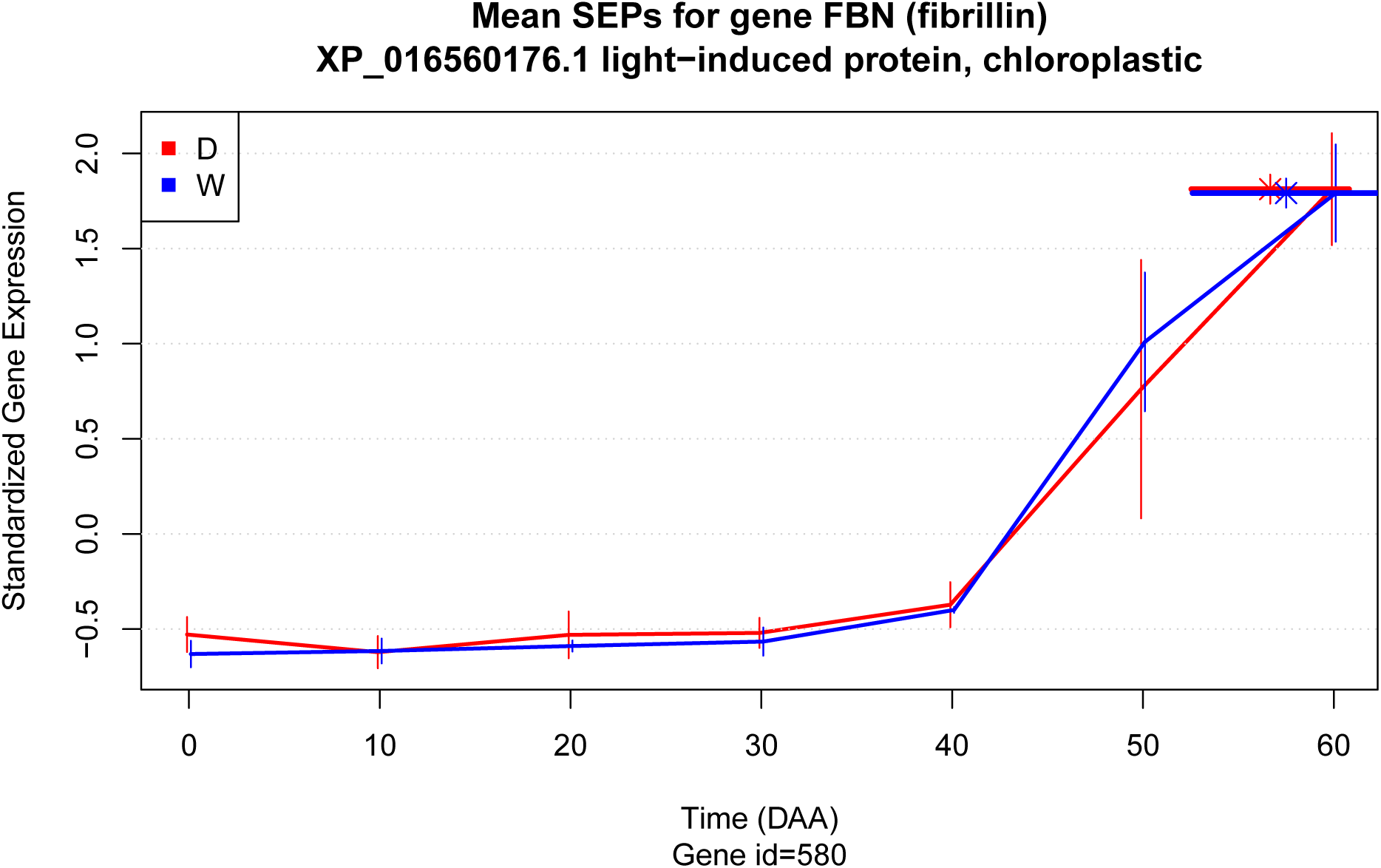
Main lines link the mean SEPs and the thin vertical lines give the 95% CI for the respective estimated points. Asterisks point to the estimated time in DAA where the maximum mean expression was estimated while broad lines over the asterisks are the 95% CI for those points.

The same plots and statistical analyses presented in figures 8 and 10 and appendices S-11 and S-11.1 for individual genes can be performed for groups of genes, as done to plot figures 2, 3 and 4 in the main text. To perform statistical analyses of a gene, or sets of genes, we considered contrasts between two groups of accessions, 6 D (AS, CW, JE, ST and ZU in Table 1) and 4 W (CO, QU, SR and SY in Table 1 in main text). In all cases, the null hypothesis was that at each time point the mean expression of the D and W groups was equal, whereas the alternative was that these parameters differed. Variation within the D and W groups was considered as a statistical error (unexplained variation) and a t-test was used to obtain Confidence Intervals (CI) for the means and to evaluate significance at each of the 7 time points sampled. We determined the mean SEPs for different gene groups in the D and W accessions (Figure 11).

The mean for the D and W groups differed significantly (Figure 11 A). At the mature flower state (0 DAA), the standardized mean expression for D was much higher than for W, implying that the average transcription activity in this state is substantially larger for the D genotypes. In the interval between 0 and 10 DAA, the mean standardized expression increased for both groups, although the rate of increase was higher for D. At 10 DAA, the mean expression for D reached a peak value, but for W the increase continued, although at a slower rate, to peak at 20 DAA. From the peak at 10 DAA, the mean expression for D decreased, at different rates, and was lower at all subsequent time points. The lowest value was seen at 60 DAA. In contrast, decreases in the mean expression for W began later, occurring from 20 up to 50 DAA, and reached a minimum of −0.27, which is smaller than the minimum for the D group, −0.25, seen at 60 DAA. The more relevant differences between mean expression profiles between D and W were seen during the intervals between 10 and 20 and 50 to 60 DAA, when the trend (i.e., slope of the regression models) was inverted such that D was decreasing while W was increasing. On the other hand, less marked differences between D and W were seen between 30 and 50 DAA when the mean standardized expression decreased nearly in parallel for both groups. The average of the time at which the maximum expression was reached in each group (marked by asterisks) was five days earlier for D than W. All observed differences were significant.

**Figure 9.**
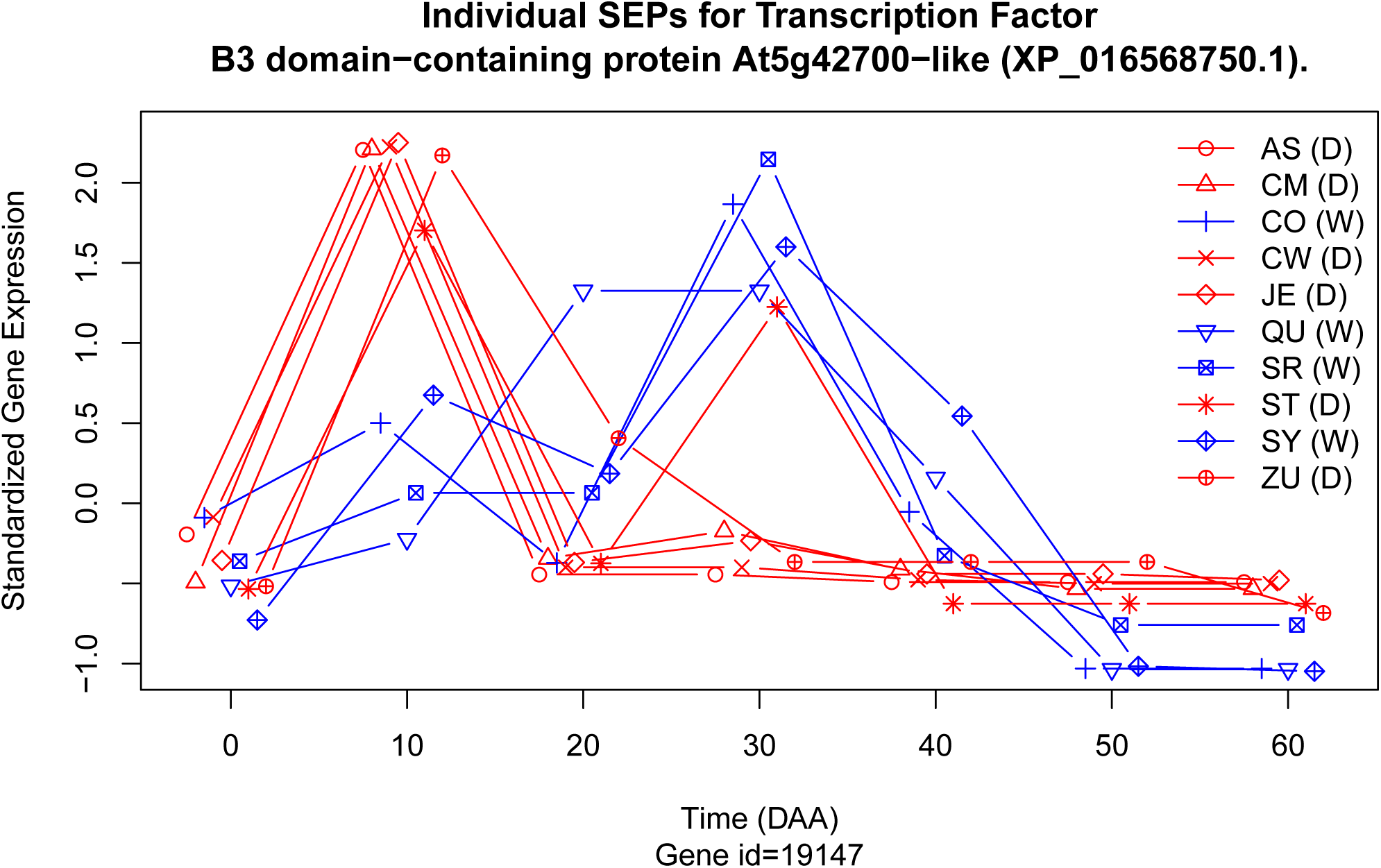
SEPs per accession for a gene with highly different expression patterns between D and W. Values in *Y* axis slightly displaced to avoid overlap.

**Figure 10.**
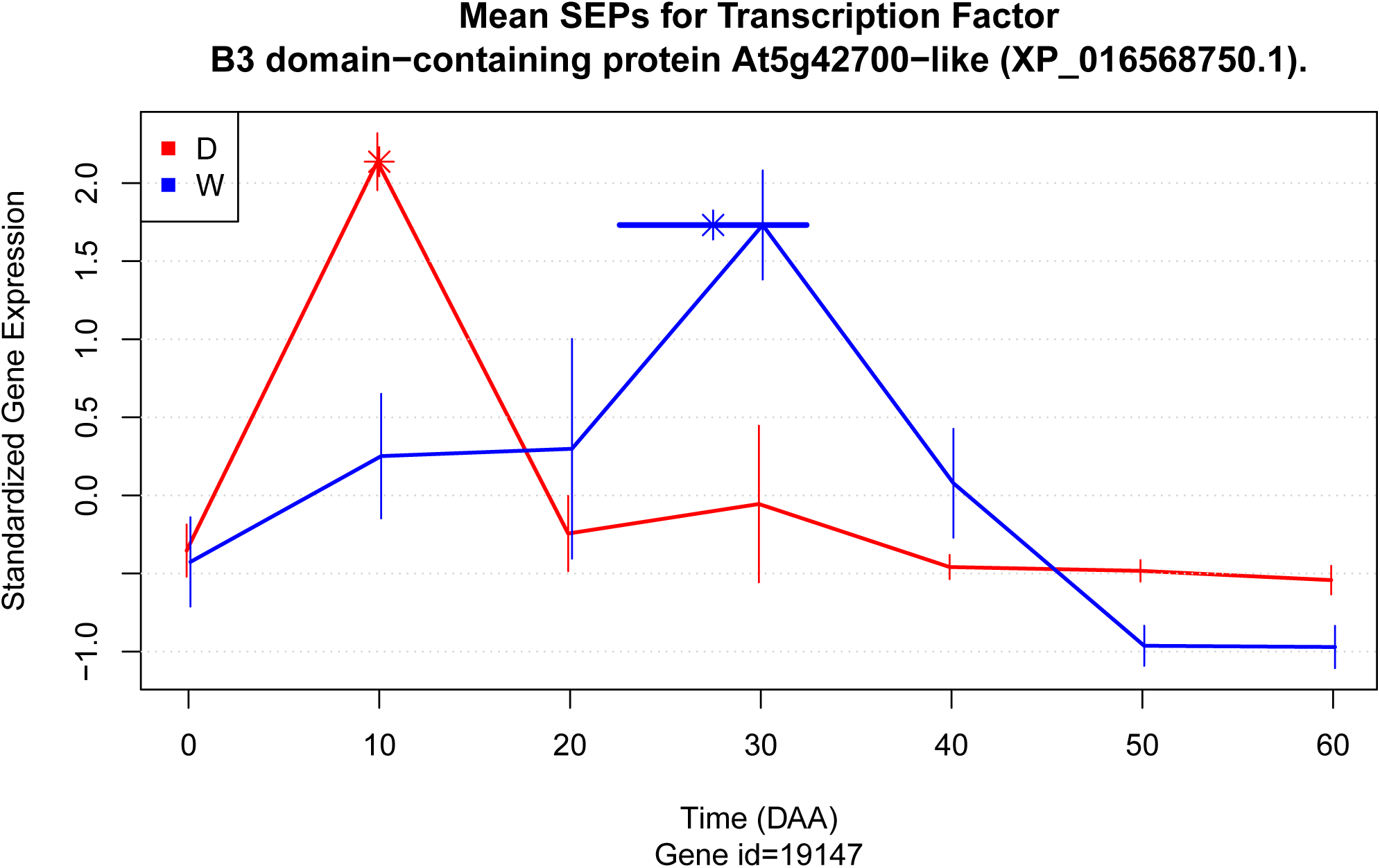
Mean SEPs per accession for a gene with highly different expression patterns between D and W.

**Figure 11.**
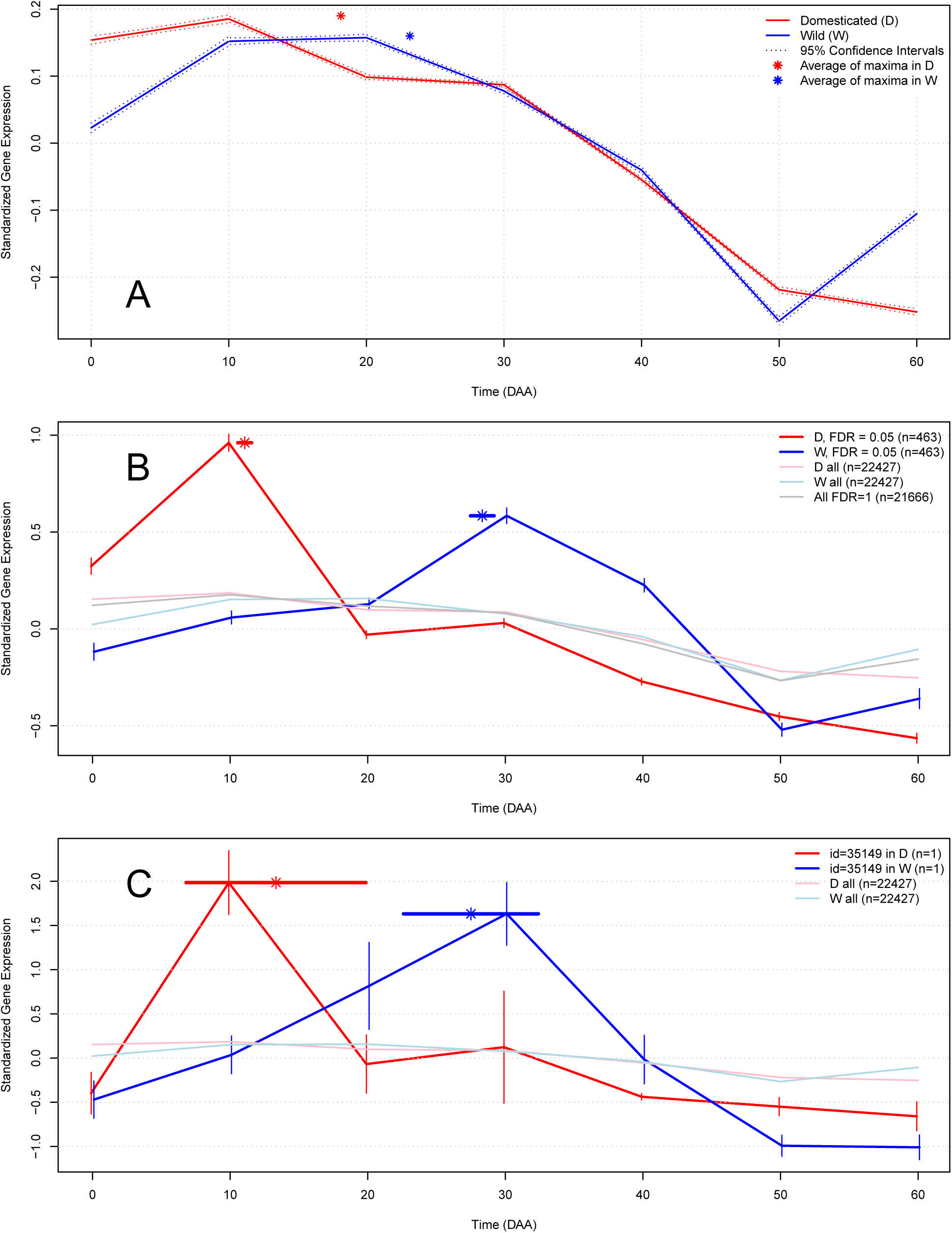
Mean SEP (Standardized Expression Profile) for groups of genes in Domesticated (D) and Wild (W) accessions. Continuous colored lines link the means of standardized gene expression at each time point. (A) Complete set of expressed genes (n=22,427). (B) Set of genes having differential expression profiles between D and W (n=463; FDR=0.05). Pale colors indicate the expression profile for all genes, and the gray line represents genes that had no difference in expression between D and W (FDR = 1). (C) Expression profiles for the gene (n=1) encoding the protein “G2/mitotic-specific cyclin S13-7” (XP 016543946.1). In B and C the thin vertical lines represent the 95% CI for the means. Asterisks indicate the mean time of maximum expression and the horizontal lines over the asterisks represent the 95% CI for the mean at each time point.

Differences in SEP of individual genes varied between D and W. To select the genes having the largest differences between D and W, we applied a statistical test on individual differences and used a False Discovery Rate (FDR) threshold of 0.05, which for these tests produced a *P* value *<* 0.000002. Using these criteria we selected a set of 463 genes, representing approximately 2.06% of the total (Figure 11 B). The expression profiles of these 463 selected genes differed markedly between D and W, ranging from −0.56 (D at 60 DAA) to 0.96 (D at 10 DAA), which is much larger than the range of variation for the means of all genes (Figure 11 B, pale red and blue lines). The profiles for these genes also completely differed from the average profile of genes that had similar expression profiles in both D and W (grey line, FDR = 1). The differences in expression profiles between D and W were well defined and significant; the peak of mean expression for D occurred at 10 DAA, while the peak for W occurred later, at 30 DAA. The average time of maximum expression (asterisks with corresponding 95% CIs) was 11.06 DAA for D and 28.33 DAA for W, or a difference of −17.27 DAA. Of the 463 selected genes, 36 (36/463 *≈* 0.08; 8%) are transcription factors (TFs). This percentage is higher than that for TFs annotated in the Capsicum genome (1,859/34,986*≈* 0.05 or 5%). A list and description of the 463 selected genes and details of statistical analyses are presented in the Supplemental SG and SM-4, respectively.

We next focused on the expression profiles in the D and W accessions for a single gene encoding the protein ‘G2/mitotic-specific cyclin S13-7’ (Figure 11 C). For this gene, the 95% confidence intervals (CIs) for the means at each time (thin vertical lines), as well as for the average of the time at which maximum expression was reached for each group (horizontal lines over the asterisks) was longer, since the means were obtained from only one gene (n=1) and thus each point is obtained from only individual data for the 6 and 4 accessions for D and W, respectively (see Methods). Nevertheless, the sample size and statistical method employed show that there are significant differences between the D and W profiles for a single gene, given that the 95% CI values do not overlap (Figure 11 C).

The results indicate that the design and results of this experiment showed differences in expression profiles between D and W at the level of whole gene sets (Figure 11 A), groups of particular genes (Figure 11B), and individual genes (Figure 11 C). Taking these findings together, we can thus conclude that there are relevant differences in expression profiles between domesticated and wild varieties of chili peppers.

#### S-4.1. Differences in Expression of Genes Related to Cell Reproduction Appear Earlier and are Larger in Domesticated than Wild Genotypes

Based on the evidence that mean SEP differ between the D and W accessions, we investigated differences in expression profiles in groups of genes related to particular biological processes. We first examined the mean SEPs of a group of 1,125 genes associated with cell reproduction (Figure 12).

We observed that the mean tendency of all 1,125 genes (solid lines) and a subset of 170 genes showed significant (*P <* 0.01) differences in expression profiles between D and W (dashed lines; Figure 12 A). Moreover, significant differences between D and W were observed at all 7 time points for both the entire group and gene subset. For both groups (n=1,125 and n=170), the mean expression was higher in D than for W at 0, 10 and 50 DAA. Meanwhile, the intervals from 10 to 20 and 50 to 60 DAA had contrasting tendencies for D and W. For both intervals the mean expression decreased for D, but increased for W. The peak of mean expression occurred earlier for D (at 10 DAA) than for W (at 30 DAA) and the magnitude of expression at the peak was also much larger for D than for W.

The mean expression value for 235 genes that are directly annotated in the cell cycle —but not in other cell reproduction processes— was significantly higher and occurred earlier for D compared to W, as evidenced by the peak of 0.3 standardized units at 10 DAA for D and 0.2 standardized units 30 DAA for W (Figure 12 B). Similarly, the mean expression for 69 kinesins or kinesin-related proteins among the 1,125 genes associated with cell reproduction exhibited a differential expression peak at 10 DAA for D accessions, but for W accessions the peak was later at 30 DAA (Figure 12 C).

**Figure 12.**
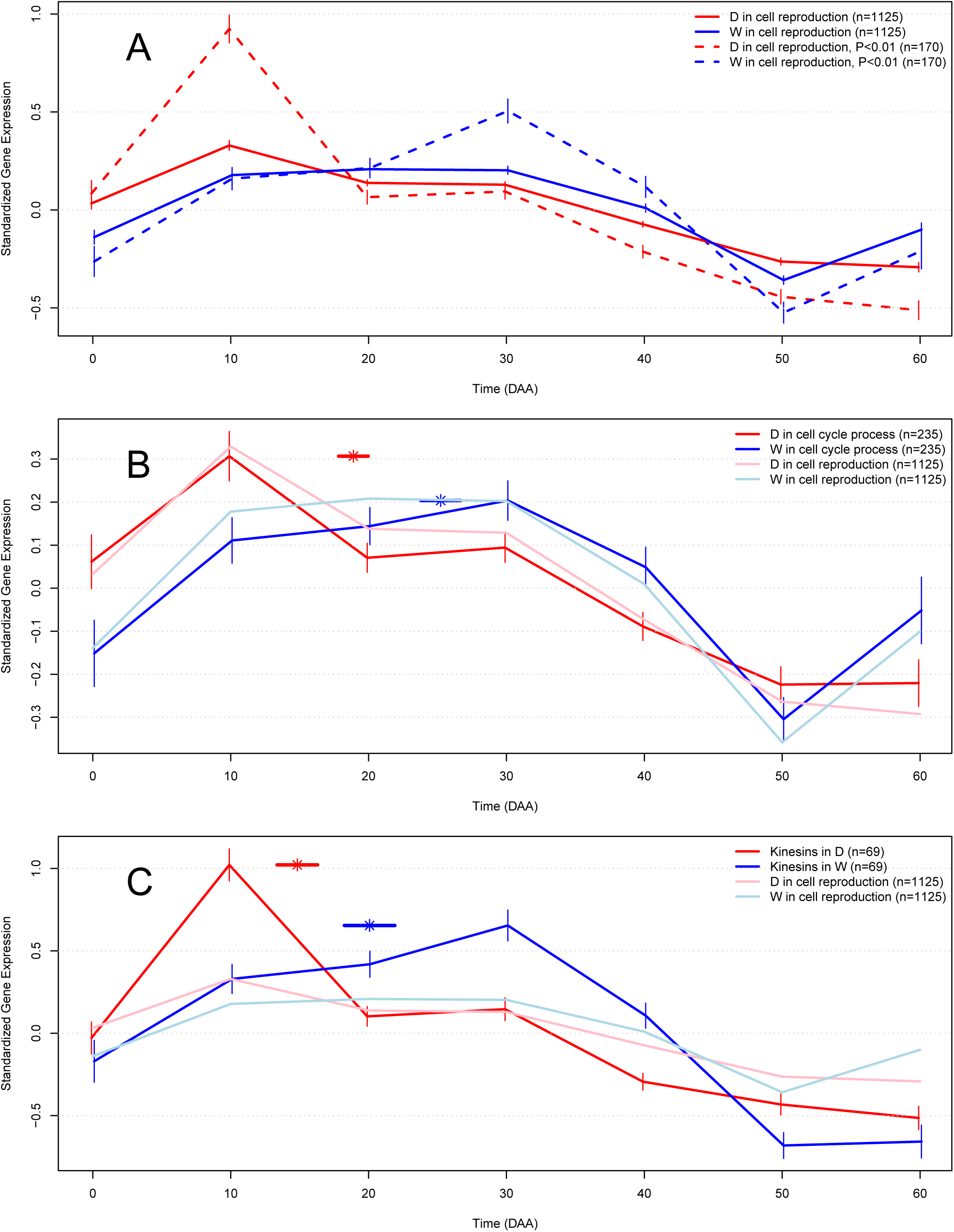
Mean Standardized Expression Profile (SEPs) for groups of genes associated with cell reproduction in Domesticated (D) and Wild (W) accessions. Vertical lines indicate 95% CI, asterisks denote mean time of maximum expression and horizontal lines over asterisks represent the 95% CI for the parameter. (A) Solid lines show the expression profile for the entire set of 1,125 genes and dashed lines represent expression of a set of 170 genes that had the highest differential expression between the D and W groups (*P <* 0.01). Genes annotated in (B) cell cycle process and (C) Kinesins.

Thus, changes in expression of genes associated with cell reproduction were significantly larger and occurred earlier for D relative to W accessions, not only for the full set of genes, but also for particular bioprocesses and gene families (Figure 12).

#### S-4.2. Biological Processes Enriched in Genes That Are Expressed Earlier in Domesticated Genotypes

The results presented above indicate that SEPs in D and W accessions undoubtedly differ (Figure 11), and genes for which expression peaks at 10 DAA for D but at 30 DAA for W (denoted here as ‘D10W30’) play an important role in cell reproduction (Figure 12). To validate and expand our study, we considered 542 genes having the D10W30 expression pattern in a Gene Ontology enrichment analysis.

A total of 86 biological processes (BPs) were significantly enriched (FDR = 0.05; *P <* 0.0015) in the D10W30 set, with a median odds ratio of 9.5. As such, these genes were much more abundant in these BPs than would be expected by chance. Apart from the abovementioned BPs related to cell reproduction, 43 of the enriched BPs, or 50% of the total, are involved in either positive or negative regulation of various biological processes. Of these, 4 (5%) are related to cellular component organization or biogenesis, 3 are associated with cellular component assembly, and another 3 play roles in organelle organization or fission. The general bioprocess “cellular process” (GO:0009987) is also highly enriched in the D10W30 gene set, with an odds estimate of 2.25 and a highly significant P-value of 2.76 *×* 10^*−*8^.

These results consider the expression patterns of sets of genes grouped by D and W accessions. Next we considered SEPs for single genes (Figure 13). For the three highlighted genes, the mean expression values for D occur at 10 DAA, while for W the means are observed at 30 DAA, consistent with the pattern D10W30. However, the expression patterns for individual accessions (dotted lines) are variable, even when the mean tendency (continuous lines) consistently followed the D10W30 pattern (Figure 13 A to C).

In examining the expression patterns for the gene encoding the “high mobility group B protein 6”, a WRKY transcription factor involved in the nucleosome/chromatin assembly that was annotated in 12 of the 86 abovementioned BPs, particularly cell reproduction BP, there are two outliers among the D10W30 pattern (Figure 13 A). Accession ST (D) had an expression peak at 30 DAA rather than at 10 DAA-even though it had a local maximum at 10 DAA. Accession SY (W) had an expression peak at 40 DAA instead of at 30 DAA. However, the average expression pattern for this gene conforms to the D10W30 pattern and the differences in mean expression between D and W are significant at the two critical points, 10 DAA and 30 DAA.

The gene encoding the transcription factor “MYB-related protein 3R-1” was included in 6 of the 86 enriched BPs and is mainly related to cellular, chromosome and organelle organization. Notably, in comparing Figures 13 A and 13 B, the same accessions, ST (D) and SY (W), are outliers among genes showing the D10W30 pattern, and both had the same tendencies, i.e., high expression at 30 DAA for ST (D) and a late peak at 40 DAA for SY (W). On the other hand, differences in mean expression between D and W were significant at the two critical points 10 DAA and 30 DAA (Figure 13 A, B).

The “kinetochore protein NDC80” is part of multiprotein kinetochore complexes that couple eukaryotic chromosomes to the mitotic spindle to ensure proper chromosome segregation. NDC80 is part of the outer kinetochore and forms a heterotetramer with proteins NUF2, SPC25 and SPC24 (Santaguida and Musacchio, 2009; D’Archivio and Wickstead, 2017). Interestingly, the genes encoding NUF2 and SPC25 also exhibit the D10W30 expression pattern. NDC80 is conspicuously present in 74 of the 86 enriched BPs (Figure 13 C).

**Figure 13.**
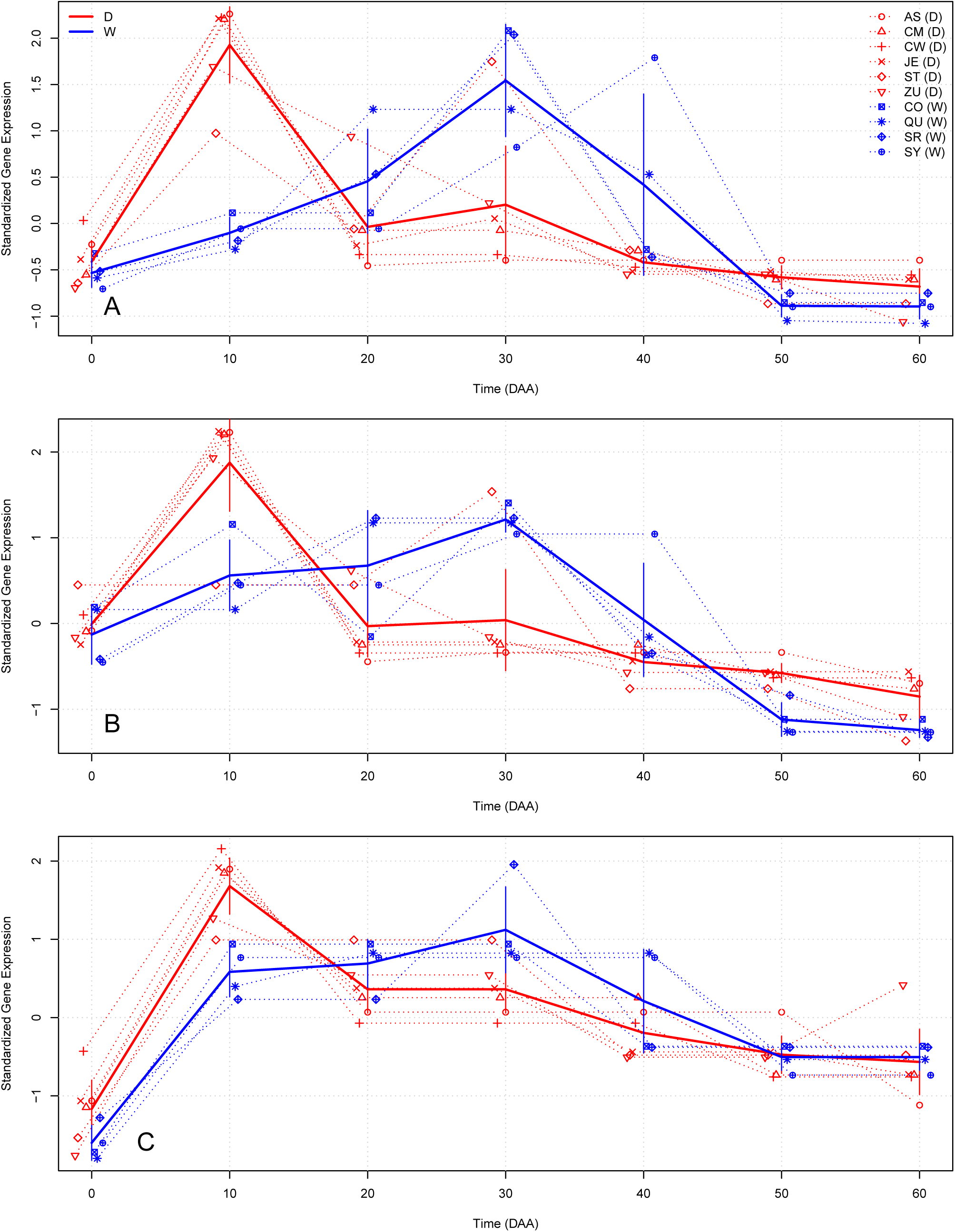
Gene expression patterns for three genes having the D10W30 expression pattern in Domesticated (D) and Wild (W) accessions. Dashed lines show the SEPs for each accession, and solid lines show mean SEPs per group (D and W). Vertical lines represent 95% CI for mean values at each time. Keys correspond to those shown in Table 1. (A) High mobility group B protein 6 (XP 016555757.1); (B) MYB-related protein 3R-1 (XP 016537977.1); (C) Kinetochore protein NDC80 (XP 016539151.1).

### S-5. Gene Ontology (GO) enrichment analyses

After discovering that mean SEP in the D accessions was different to the one in the W group (Figure 1A in the main text), we confronted the problem of finding the functional meaning of that difference, and for this aim we employed Gene Ontology or ‘GO’ annotations (Ashburner et al., 2000). We isolated the set of genes with a more extreme difference, the *n* = 463 genes with a False Discovery Rate, FDR = 0.05 (Benjamini and Hochberg, 1995), presented in Figure 1B of the main text, and noticed that this group presented the pattern ‘D10W30’, where the maximum mean expression was at 10 DAA in D, while such maximum occurred at 30 DAA in W (Figure 1B in the main text). Furthermore, we found that a set of 542 genes presented SEPs with D10W30 patterns, and this set was one of the targets for GO enrichment analyses, employing the ‘Biological Process’ GO ontology and motivated by the results in (Lægreid et al., 2003).

To perform GO enrichment analyses we considered the total population of 22427 genes expressed during fruit development of which 12102 are annotated in one or more of the 2547 GO biological processes annotated in chili. We are interested in the property of a gene to belong to a specific GO category, with the aim to establish whether the class of genes with a specific expression pattern, e.g. genes with mean SEPs D10W30, presented an enrichment in the GO Biological Process of interest with respect to the total gene population. Among the different tests that could be used to test association between a target gene set and a functional GO Biological Process (Rivals et al., 2007), we selected the Fisher’s exact test.

We programed a function to summarize the results of the test, and employing different targets performed the analyses of the 2547 GO biological processes, evaluating the *P*-value of each result, and transforming it to a *Q*-value to have a FDR (Benjamini and Hochberg, 1995) of 5%. To take into account the structure of the GO ontology, which is fundamental to the analyses interpretation (Rhee et al., 2008), we performed a filtering of redundant and highly correlated biological process using a gene network approach.

As an example of the analyses performed, Appendix S-11.2 presents the R output for the ‘Cell Cycle’ biological process having as target the set of 542 genes with D10W30 patterns. In Appendix S-11.2 we can see the output of function ‘BP.analysis.ById’. This function gives the observed and expected 2 *×*2 contingency tables as well as the full results of Fisher’s exact test, making easier result’s interpretation.

Sheet ‘Bio Process’ in the excel file “SG.xlsx” of ‘Supplemental Information’ presents the full results of the analyses of the 2547 GO biological processes using as target the set of genes with pattern D10W30.

### S-6. Genes and Bio Processes (BPs) reported

Excel file “SG.xlsx” in ‘Supplemental Information’ includes four sheets with the following results:

**Gene :** Data for the 22427 genes expressed during fruit development (in table “gene” of the SALSA database).

**Gene column definitions :** Column definitions for the “Gene” sheet.

**id:** Numerical identifier in the SALSA database.

**ProtId:** Protein identifier of the gene product (if known, otherwise NULL).

**Prot Desc:** Protein short description (if known, otherwise NULL).

**URL:** URL for UniProtKB database using Prot Desc (if known, otherwise NULL).

**isTF:** Is the gene product annotated as Transcription Factor? (T if True, F if False).

**D10W30:** Is the SEP of the gene of class ‘D10W30’ [see main text] (TRUE or FALSE).

**BioProc:** Is the gene product annotated in one or more GO Bio Processes (T if True, F if False).

**ZunlaDom:** Is the gene annotated with domestication footprint in Qin et al. (2014)? NULL if it is not annotated as such, otherwise the name of the gene reported by Qin et al. (2014) is given.

**P value:** P value for the test of differences of SEPs between domesticated (D) and wild (W) accessions. See main Methods and Supplemental SI-1.3.

**Q value:** P value transformed to Q value using R function p.adjust() with method = “fdr” to calculate False Discovery Rate (DFR).

**Gene id:** Genomic identifier of the gene.

**chromosome:** Chromosome where the gene is located; “NULL” if unknown see “scaffold” below.

**scaffold:** scaffold Scaffold where the gene was located (If Chromosome “NULL”).

**Strand:** Strand coding for the gene (“+” or ‘-”)

**start:** Genomic coordinate where the gene starts.

**end:** Genomic coordinate where the gene ends.

**length:** Length of the gene in base pairs (bps).

**sequence:** Gene sequence.

**Bio Process :** Data for the 2547 Gene Ontology (GO) biological processes analyzed (in table “ResBioProcess” of the SALSA database).

**Bio Process column definitions :** Column definitions for the “Bio Process” sheet.

**BP**.**id:** Numerical identifier of the Biological Process in the SALSA database.

**bio**.**process:** Gene Ontology (GO) Biological Process.

**odds:** Estimated odds in the contingency table.

**P:** P-value of the Fisher’s exact test for the 2 *×* 2 contingency table.

**AnnTarg:** Number of genes in the process which are annotated in the target.

**NotAnnTarg:** Number of genes in the process which are NOT annotated in the target.

**AnnNotTarg:** Number of genes in the process which are annotated but NOT in the target.

**NotAnnNotTarg:** Number of genes in the process which are NOT annotated and NOT in the target.

**Q:** P value transformed to Q value using R function p.adjust() with method = “fdr” to calculate False Discovery Rate (DFR).

Information in the “**Gene**” sheet was obtained from the data send by NovoGene after RNA-Seq sequencing and analyses and corresponds to the annotation in the reference genome CM334 v1.6. On the other hand, information in the “**Bio Process**” sheet was the results of the GO enrichment analyses described here in section S-5.

### S-7. Network estimation

As mentioned in (Allocco et al., 2004),

> “It is axiomatic in functional genomics that genes with similar mRNA expression profiles are likely to be regulated via the same mechanisms. This hypothesis is the basis for almost all attempts to use mRNA expression data from microarray experiments to discover regulatory networks.”

Ideally we would like to estimate a Gene Regulatory Network (GRN) for the whole chili transcriptome. That aim is practically impossible with the current incomplete knowledge of the interactions between genes in the *Capsicum* transcriptome. However, an attainable and relevant goal within the framework of our study is to estimate robust networks of functionally related genes, as the one presented in Figures 3 and 4 of the main text. Here we detail the method employed to obtain that network.

We have a total of 22,427 genes consistently expressed in all accessions, of which 352 are annotated in the BP ‘Cell Cycle’ and of these 25 belong to the class ‘D10W30’, i.e., these 25 genes present a maximum expression at 10 DAA in the 6 domesticated (D), while the maximum expression is at 30 DAA in the 4 wild (W) accessions. After examining the Euclidean distances between the SEPs of the 25 genes, we selected 6 of them which present a highly consistent SEPs in both, D and W expression. We selected the 6 structural genes presented in the network of Figure 3 and 4 of the main text (represented by orange circles in that figure) by setting a threshold of Euclidean distance 1 *≤*between pairs of gene SEPs. Table 2 presents the medians of the Pearson correlation 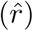 and *P* values for SEPs of the 6 Structural genes included in the network.

**Table 2.** Median Pearson Correlation 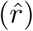 and *P* values for SEPs of the 6 Structural genes included in the network presented in Figures 3 and 4 of the main text.

|  | Between D and W | Within D | Within W |
| --- | --- | --- | --- |
| $n$ | 144 | 90 | 36 |
| $\hat{r}$ | 0.37477 | 0.92227 | 0.77658 |
| $P$-value | 0.40749 | 0.00310 | 0.04001 |
|  | Between | Within |  |
| $n$ | 144 | 126 | |
| $\hat{r}$ | 0.37477 | 0.88496 | |
| $P$-value | 0.40749 | 0.00810 | |

In Table 2 column ‘Between D and W’ presents cases where correlation was estimated for the same gene but taking one D and one W accession, thus correlations are between SEPs in D and W. The number of such pairs of different correlations equals 6 D*×* 4 W*×* 6 genes, *n* = 6 *×*4 *×*6 = 144. On the other hand, columns ‘Within D’ and ‘Within W’ present cases where correlation was estimated for the same gene but taking different accessions within the same group (D or W, respectively). The number of possible comparisons are *n* = (6*×* (6 *−* 1))*/*2*×* 6 = 90 for the column ‘Within D’ and *n* = (4 *×* (4*−* 1))*/*2*×*6 = 36 for the column ‘Within W’. In Table 2 we can see that the median of the correlations for SEPs within the D and W groups are high, 0.92227 and 0.77658 and significant (*P*-values of 0.00310 and 0.04001), respectively, while the median of the correlation for SEPs between the D and W groups was smaller, 0.37477, and not significant (*P*-value of 0.40749). Last rows of Table 2 groups columns ‘Within D’ and ‘Within W’ into a single column, ‘Within’, and from such grouping we obtain the same conclusion than above, i.e., the 6 structural genes have highly and significantly correlated SEPs within but not between accession groups.

Results in Table 2 refer to all posible pairs of the 6 structural genes. However, not all pairs of structural genes are linked (by double headed arrows) in the network of Figures 3 and 4; the genes considered as linked in the network present a value of 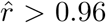 within D and W groups, with a *P*-value *<* 0.0001. In contrast the same pairs of structural genes present a value of 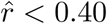 between D and W groups, with a non-significant *P*-value *>* 0.5 Thus the network of structural genes presented in Figures 3 and 4 presents a set of cell cycle genes which are highly coordinated in time within the D and W groups, presenting the expression pattern D10W30.

To corroborate that the expression of the 14 genes included into the network are indeed very well segregated, we performed a Principal Component Analysis (PCA) of the 14 *×* 10 = 140 7-dimensional SEPs, which tendency by group is presented in Figure 3 B in the main text. In the biplot shown in Figure 14 we see the expression of the 14 genes labeled by their accession of origin and with colors denoting the set of origin (D in red and W in blue), plot in the 2 dimensional space of the first 2 principal components, of which the first (*Y* axis) explains 57.7% and the second (*X* axis) explains 17.4% of the data variance, thus together the two first principal components explain 75.1% of the total variance. In this figure we can see that the first principal component efficiently segregates the data into two groups, D in the upper and W in the lower parts of the plot, with only a few outliers. Observing the eigenvectors (dark red arrows in the plot, labeled by the time DAA: 0, 10, *…*, 60) we see that the one at time 0 is almost horizontal, and thus have very small influence in the coordinates transformation. This makes sense, because at 0 DAA both groups (D and W) share the same level of expression, as shown in Figure 3 B in the main text. In contrast all the other 6 eigenvectors, corresponding to times 10, 20, *…*, 60 DAA, have a strong influence in the segregation of the D and W sets, as previously observed in Figure 3 B in the main text.

**Figure 14.**
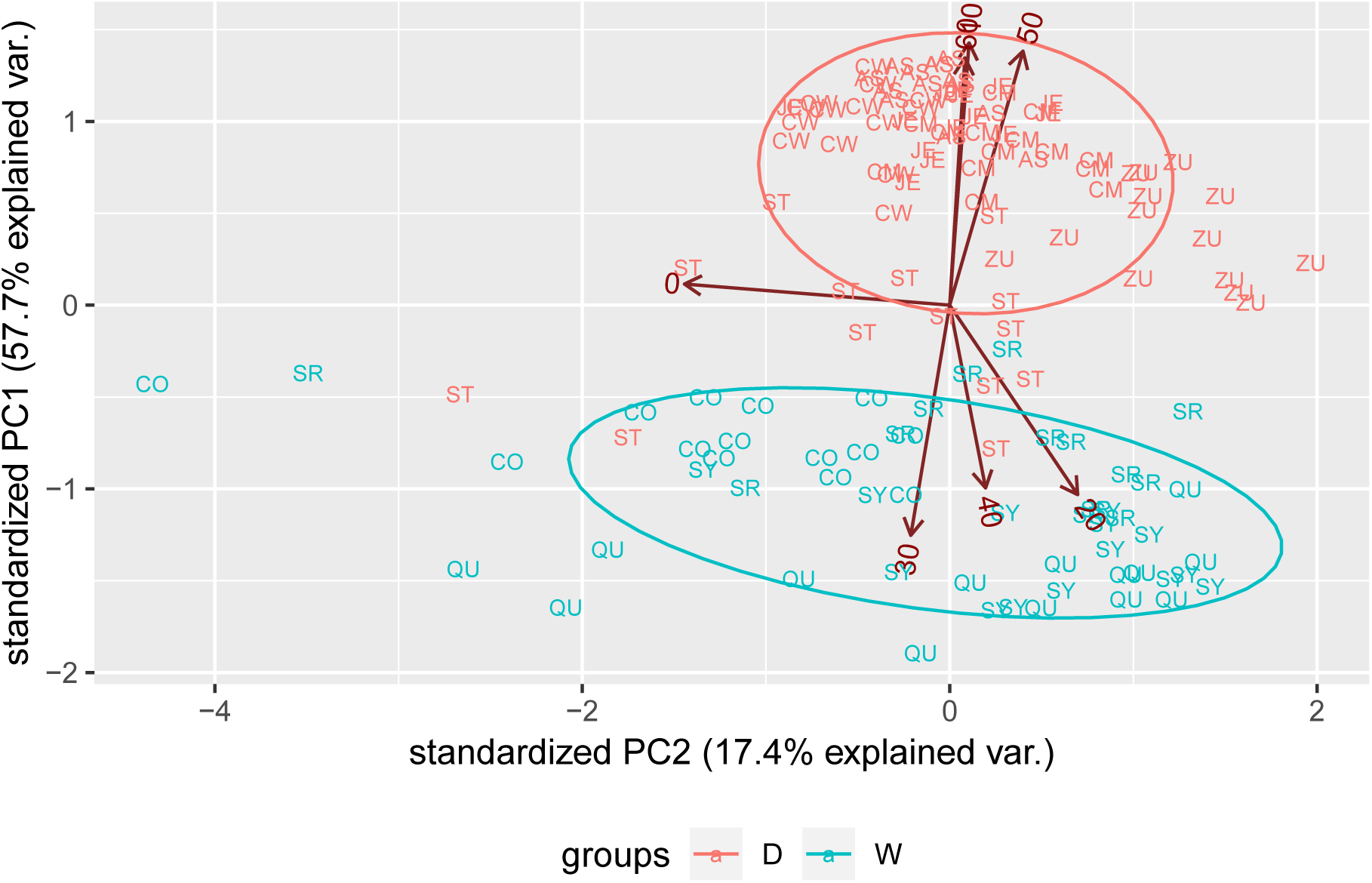
PCA analysis of the SEPs for the 14 genes in the network (Figure 2 B).

### S-8. Transcription Factor (TF) imputation

The authors of Allocco et al. (2004) analyzed 611 microarrays and found that the correlation between expression profiles of two genes must be larger than *r≈* 0.84 to have a 50% chance of sharing a common transcription factor binder. Here we assume that a target gene and a TF which is regulating it will share very alike expression patterns (SEPs) and developed an statistical approach to select a set of candidate TFs. This approach was implemented in an R function which performs the following steps:

#### Algorithm for TF imputation

1. Basic input: id - Identifier of the target gene; min.r - Threshold for the minimum correlation value, *r >* 0; min.rOm.a - Threshold for the minimum ratio of *r/ma*, where *ma* is the maximum of the absolute difference between the SEPs of the target gene and the SEP of a TF; acc.set - The set of accessions where the search will be performed.
2. Obtains from the database all SEPs for all TFs in all accessions that belong to acc.set.
3. Obtains from the database all SEPs for target gene (id) in all accessions that belong to acc.set.
4. For each accession that belong to acc.set calculates the correlation, *r*, and *r/ma* between the SEPs of each TF and the target gene. Keeps only the cases where *r≥* min.r AND *r/ma≥* min.rOm.a.
5. Obtains a final list of candidate TFs by founding the intersection of all the sets of candidates in each one of the accessions defined int the input (acc.set).
6. Output the list of TFs candidates (if any) as well as variables to judge the adequacy of each TF candidate.

It is important to consider two facts about the above described method. Firstly, parameters *r ≥* min.r AND *r/ma ≥* min.rOm.a are selected in an ‘*per accession*’ base; i.e., they are compared only with the SEPs of the TFs in the same accession. Assume that a given target gene, id, is regulated by the same TF, say, x, but that target gene has very different expression pattern in two different accessions. If id is regulated by x in both accessions, the method will likely report x in both accessions at step (4), and thus x will be part of the final output in (6). Secondly, and more important, given that the data in all accessions are fully independent, the probability of reporting ‘erroneous’ or ‘spurious’ TFs decreases exponentially with the number of accessions taken into account. This is, if the probability of reporting a spurious TF in any of the *k* accessions is *ε*, then the probability that the procedure reports the same spurious transcription factor in *k* accessions is *ε*^*k*^, e.g. if *ε* = 0.5 and *k* = 6 we have *ε*^*k*^ = 0.5^6^*≈* 0.016 and if *k* = 10, *ε*^*k*^ = 0.5^10^*≈* 0.001, etc. Under the null hypothesis of no true correlation between two arbitrary SEPs, the true value of the correlation parameter, *ρ*, is uniformly distributed in the interval [*−*1, 1], and if we restrict ourselves to positive values, *ρ≥* 0, the parameter space is simply [0, 1], and by setting a threshold min.r = 1*− ε* and employing *k* independent accessions in the determination we can effectively fix any desired error probability to be (1*− ε*)^*k*^. Furthermore, by additionally asking that *r/ma≥* min.rOm.a we will filter cases where the correlation, *r*, is high but at the same time there is an outlier in one of the times, where the maximum of the absolute value, *ma*, is large. This additional filter adds stringency to the selection method.

After running the algorithm to estimate the TF candidates for each one of the structural genes, we found the 8 TFs which are shown in Figure 3 A as blue circles and in rows 7 to 14 in Table 2 of the main text. The algorithm was run with parameters min.r = 0.5, min.rOm.a = 0.9 with the full set of 10 accessions. The next box presents the summaries of auxiliar estimates that help to calculate the robustness of the TF candidates.

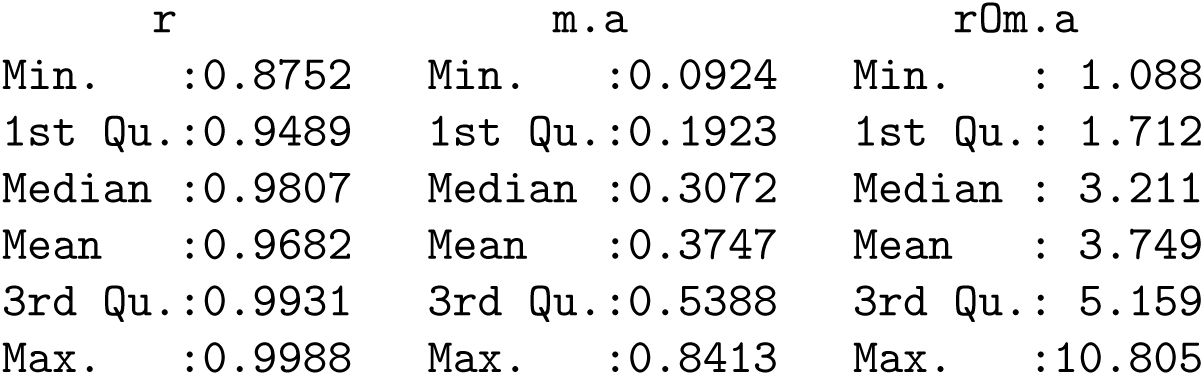

The box above summarizes the results for the 8 TFs selected, which are potential regulator of 3 of the structural genes, as shown in Figure 3 A in the main text. The statistics shown are produced from the estimation of 10 *×* (4 + 4 + 1) = 90 cases, that arise because each one of the 3 TFs was evaluated in 10 accessions, and two of them are potential regulators of 4 structural genes and one of them is potential regulator of 1 gene. By taking the mean of the 90 *r* values, 0.9682, the realized error probability is estimated as 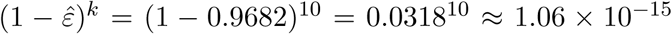, a vanishing small quantity, thus we can be reasonably sure that the relations found between the structural genes and TFs are, at least for some of the cases, very likely to reflect either, direct or indirect regulation of structural genes by the TF candidates.

The algorithm presented in this section for TF imputation was applied in our data to nominate TF candidates for the AT3 gene, resulting in the selection of only two TF, precisely the ones that have been experimentally validated as regulators of AT3 (Arce-Rodríguez and Ochoa-Alejo, 2017; Zhu et al., 2019; Sun et al., 2019). The fact that our approach recovers experimentally validated TFs demonstrates that this approach retrieves strong TFs candidates.

### S-9. Supplementary descriptions and web links for genes in the network

#### Descriptions

Items in this list give a short description and references for genes in the network of Figures 3 A and 4 and Table 2 in the main text. In each case the *Capsicum* protein identifier from Table 3 is followed by the putative *Arabidopsis* ortholog between parenthesis. Order in this list is the same than the one presented in Table 2 of the main text as well as in the rows of tables 3 and 4 presented below.

1. XP 016564755.1 (AT5G51600) 65-kDa microtubule-associated protein 3 (MAP65/ASE1). Members of the AtMAP65 family –to which AT5G51600 belongs, link membrane and microtubule dynamics during plant cytokinesis, the part of the cell division process during which the cytoplasm of a single cell divides into two daughter cells. It appears that these proteins are required to coordinate cytokinesis with the nuclear division cycle, and some MAP65 family members are known to be targets of cell cycle-regulated kinases (Steiner et al., 2016).
2. XP 016538322.1 (AT2G44190) QWRF motif-containing protein 6 (DUF566). It has been demonstrated that ENDOSPERM DEFECTIVE1 (EDE1), a mutant of the AT2G44190 gene, is expressed in the endosperm and embryo of developing seeds, and its expression is tightly regulated during cell cycle progression (Pignocchi et al., 2009). Furthermore, the authors show that EDE1 protein accumulates in nuclear caps in premitotic cells, colocalizes along microtubules of the spindle and phragmoplast, and binds microtubules in vitro. The aforementioned paper concludes that this gene codes for a microtubule-associated protein (DUF566), essential for seed development in *Arabidopsis*.
3. XP 016541615.1 (AT4G21270) Kinesin 3 isoform X3 (kinesin 1). The spindle is critical for chromosome segregation, and kinesins play crucial roles in spindle structure; in particular the *Arabidopsis* ATK1 gene (AT4G21270) is required for spindle morphogenesis in male meiosis (Chen et al., 2002). Even when XP 016541615.1 is identified as kinesin 3 (row 3 in Table 3), it is more alike with the kinesin 1 of *Arabidopsis* (alignments obtained by blastx in Appendix S-11.1) and thus it is identified with AT4G21270 in Table 4.
4. XP 016575449.1 (AT5G51600); see item (1) in this list and (Steiner et al., 2016).
5. XP 016577799.1 (AT4G20900) Protein POLLENLESS 3 (TPR). Members of the tetratricopeptide repeat (TPR) superfamily had been found in cell cycle clusters during apple fruit development (Janssen et al., 2008) and it had been demonstrated that their expression is highly regulated in early developing that fruit (Soria-Guerra et al., 2011).
6. XP 016548908.1 (AT3G44960) Shugoshin. Shugoshin protects the sister chromatid cohesion complex (cohesin) for proper chromosome segregation in mitosis, until kinetochores are properly captured by the spindle microtubules (Kitajima et al., 2006)
7. XP 016568750.1 (AT5G42700) B3 domain protein (AP2/B3-like transcriptional factor family protein) The plant-specific B3 superfamily includes families, such as the auxin response factor (ARF) family and the LAV family, as well as less well understood families, such as RAV and REM. There are indications that the B3 domain evolved on the plant lineage before multicellularity (Swaminathan et al., 2008), and, for example, the over-expression of an *Arabidopsis* B3 TF, ABS2/NGAL1 leads to the loss of flower petals (Shao et al., 2012).
8. XP 016555757.1 (AT4G11080) High mobility group B protein 6 (HMG). The high mobility group B protein 6, belongs to the HMG (high mobility group) box proteins, which is a group of chromosomal proteins that are involved in the regulation of DNA-dependent processes such as transcription, replication, recombination, and DNA repair (Johns, 2012).
9. XP 016543946.1 (AT3G11520) G2/mitotic-specific cyclin S13-7 (CYCLIN B1;3) is a regulatory protein involved in mitosis and, importantly, it is first activated in the cytoplasm and that centrosomes may function as sites of integration for the proteins that trigger mitosis (Jackman et al., 2003).
10. XP 016547461.1 (AT1G26760) B3 domain-containing protein (SET domain protein 35). AT1G26760 received high scores for plastids localization (Schwacke et al., 2007), and has been also reported in maintaining H3K4 methylation (Liu and Gong, 2011).
11. XP 016575946.1 (AT5G58280) B3 domain-containing protein At5g58280; AP2/B3-like transcriptional factor family protein. This gene has been reported to be differentially expressed in the flower and seed in *Brassica rapa*, castor bean, cocoa, soybean, and maize (Peng and Weselake, 2013), with tissues of preferential expression of the orthologous B3 gene pairs in Arabidopsis and rice.
12. XP 016574880.1 (AT1G34355) FHA domain-containing protein PS1; forkhead-associated (FHA) domain-containing protein. An insertional mutation of AT1G34355, the AtPS1 gene has been characterized and found to lead to the production of diploid pollen grains (d’Erfurth et al., 2008).
13. XP 016537977.1 (AT4G32730) Myb-related protein 3R-1 (Homeodomain-like protein). In plants, this class of Myb proteins are believed to regulate the transcription of G2/M phase-specific genes; in particular MYB3R1 act as transcriptional activator and positively regulate cytokinesis. In addition, MYB3R1 may play an important role during fruit development by regulating G2/M-specific genes (Haga et al., 2011).
14. XP 016565918.1 (AT3G22780). Protein tesmin/TSO1 CXC 3; Tesmin/TSO1-like CXC domain-containing protein. TSO1 is a protein that modulates cytokinesis and cell expansion in *Arabidopsis* (Hauser et al., 2000).

**Table 3.** NCBI links and descriptions of genes in Figure 3 A and Table 2 in the main text.

| Row | id | Prot. Id (link) | Short Protein Description |
| --- | --- | --- | --- |
| 1 | 673 | XP_016564755.1 | 65-kDa microtubule-associated protein 3 |
| 2 | 6090 | XP_016538322.1 | QWRF motif-containing protein 6 |
| 3 | 15446 | XP_016541615.1 | kinesin 3 isoform X3 |
| 4 | 19658 | XP_016575449.1 | 65-kDa microtubule-associated protein 3 isoform X1 |
| 5 | 19813 | XP_016577799.1 | protein POLLENLESS 3 |
| 6 | 24546 | XP_016548908.1 | shugoshin-1; meiotic chromosome segregation |
| 7 | 19147 | XP_016568750.1 | B3 domain Prot. At5g42700 |
| 8 | 24186 | XP_016555757.1 | high mobility group B protein 6 |
| 9 | 35149 | XP_016543946.1 | G2/mitotic-specific cyclin S13-7 |
| 10 | 5824 | XP_016547461.1 | SET domain; methyltransferase activity; LOC107847605 |
| 11 | 11410 | XP_016575946.1 | B3 domain-containing protein At5g58280 |
| 12 | 12656 | XP_016574880.1 | FHA domain-containing protein PS1 |
| 13 | 13605 | XP_016537977.1 | Myb-related protein 3R-1 |
| 14 | 7175 | XP_016565918.1 | protein tesmin/TSO1 CXC 3 |

**Table 4.**
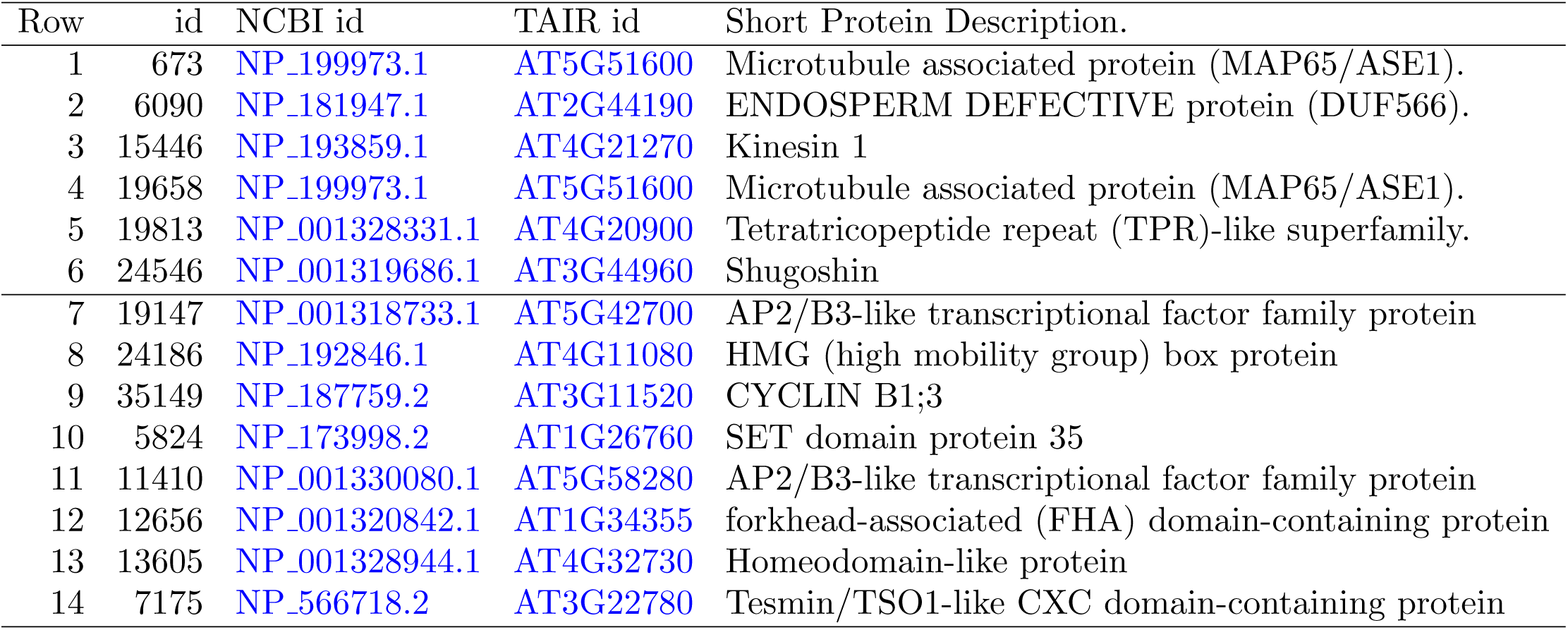
Putative *Arabidopsis* orthologous of genes in Figure 3 A and Table 2 in the main text.

| Row | id | NCBI id | TAIR id | Short Protein Description. |
| --- | --- | --- | --- | --- |
| 1 | 673 | NP_199973.1 | AT5G51600 | Microtubule associated protein (MAP65/ASE1). |
| 2 | 6090 | NP_181947.1 | AT2G44190 | ENDOSPERM DEFECTIVE protein (DUF566). |
| 3 | 15446 | NP_193859.1 | AT4G21270 | Kinesin 1 |
| 4 | 19658 | NP_199973.1 | AT5G51600 | Microtubule associated protein (MAP65/ASE1). |
| 5 | 19813 | NP_001328331.1 | AT4G20900 | Tetratricopeptide repeat (TPR)-like superfamily. |
| 6 | 24546 | NP_001319686.1 | AT3G44960 | Shugoshin |
| 7 | 19147 | NP_001318733.1 | AT5G42700 | AP2/B3-like transcriptional factor family protein |
| 8 | 24186 | NP_192846.1 | AT4G11080 | HMG (high mobility group) box protein |
| 9 | 35149 | NP_187759.2 | AT3G11520 | CYCLIN B1;3 |
| 10 | 5824 | NP_173998.2 | AT1G26760 | SET domain protein 35 |
| 11 | 11410 | NP_001330080.1 | AT5G58280 | AP2/B3-like transcriptional factor family protein |
| 12 | 12656 | NP_001320842.1 | AT1G34355 | forkhead-associated (FHA) domain-containing protein |
| 13 | 13605 | NP_001328944.1 | AT4G32730 | Homeodomain-like protein |
| 14 | 7175 | NP_566718.2 | AT3G22780 | Tesmin/TSO1-like CXC domain-containing protein |

## S-10. Appendix (R output)

### S-11. Analyses of gene with id=580 (FBN); see Figure 8 which presents the plot obtained with the function

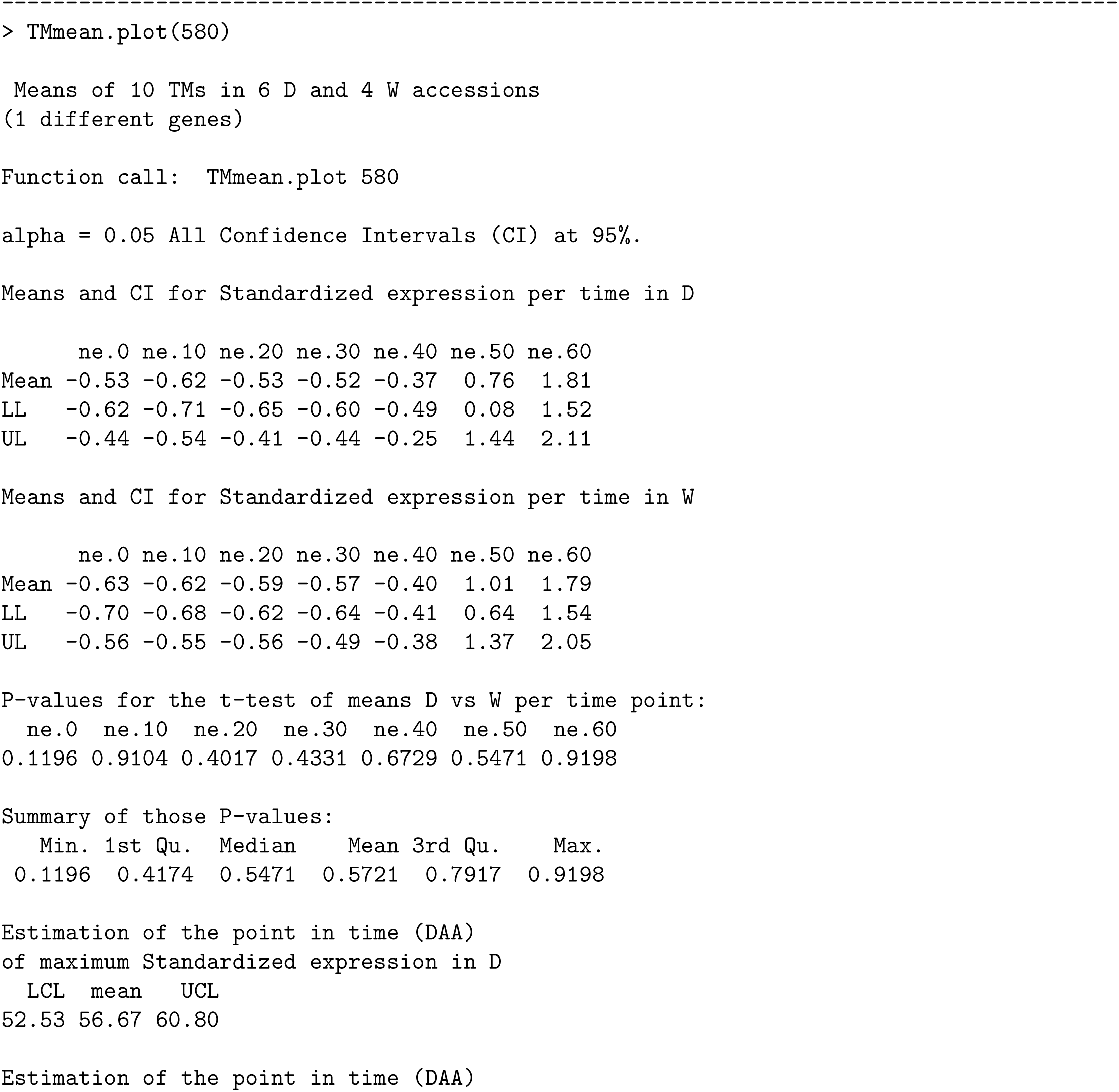
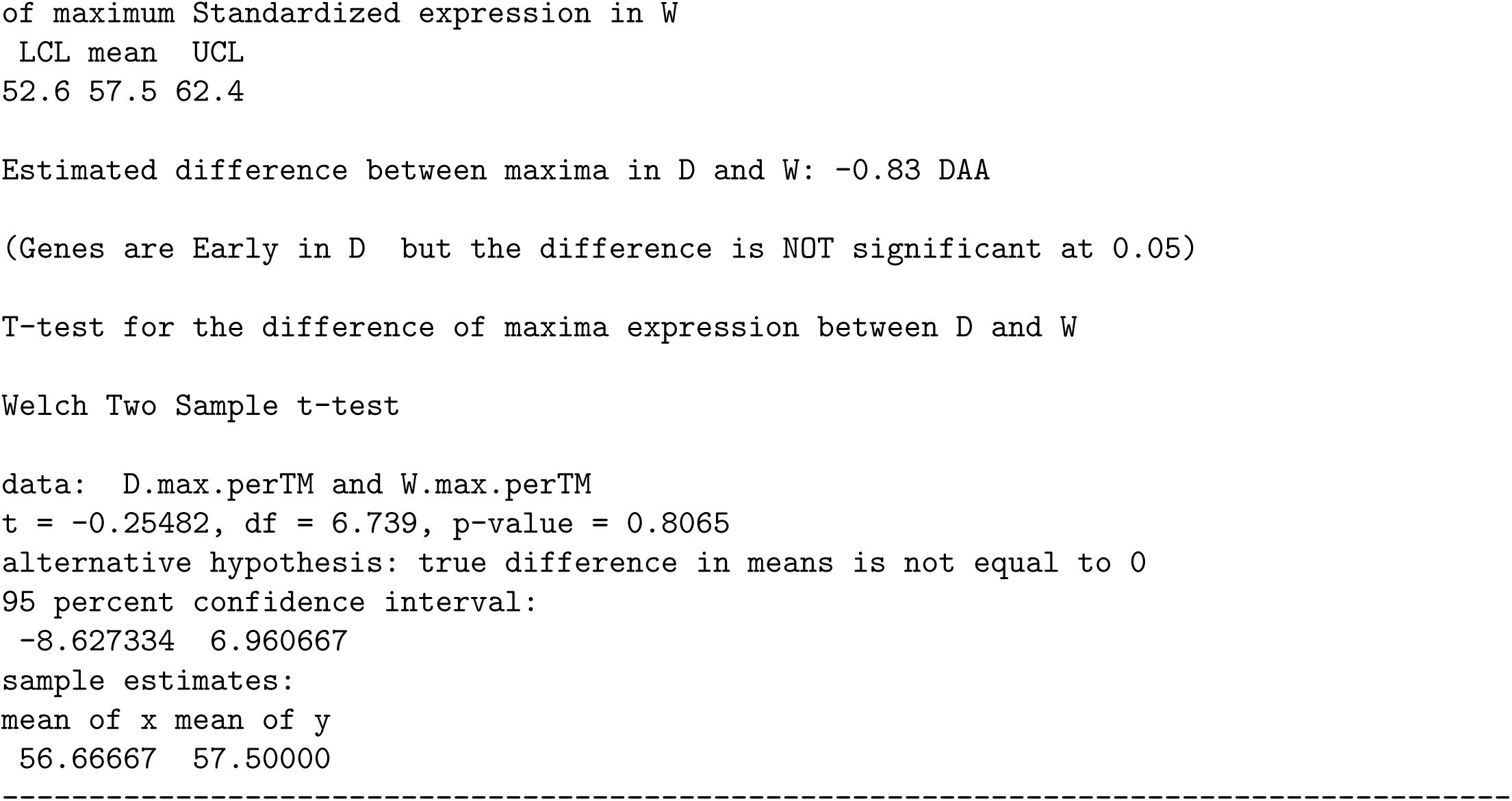

#### S-11.1. Analyses of gene with id= 19147 (B3 domain-containing protein); see Figure 10 which presents the plot obtained with the function

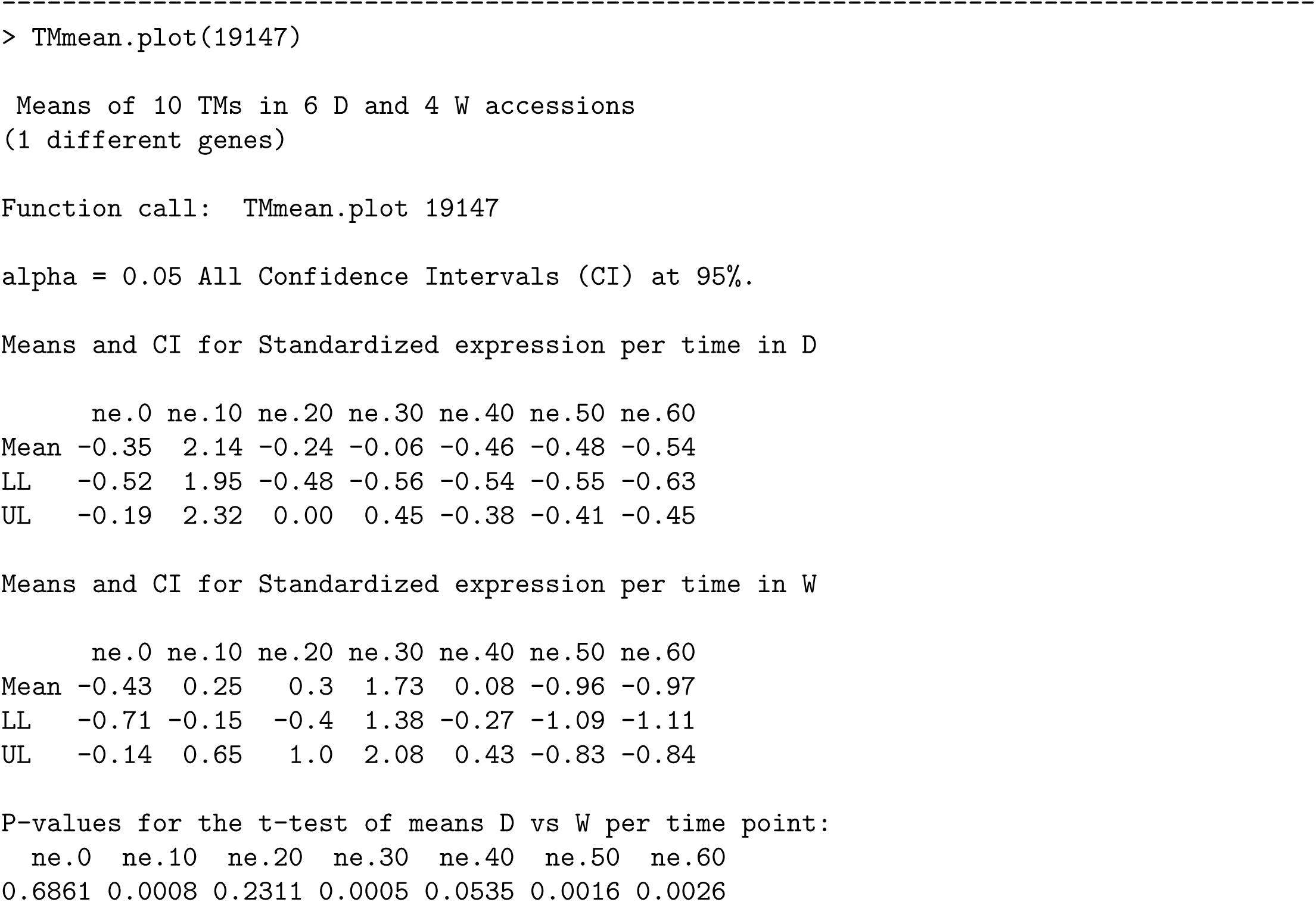
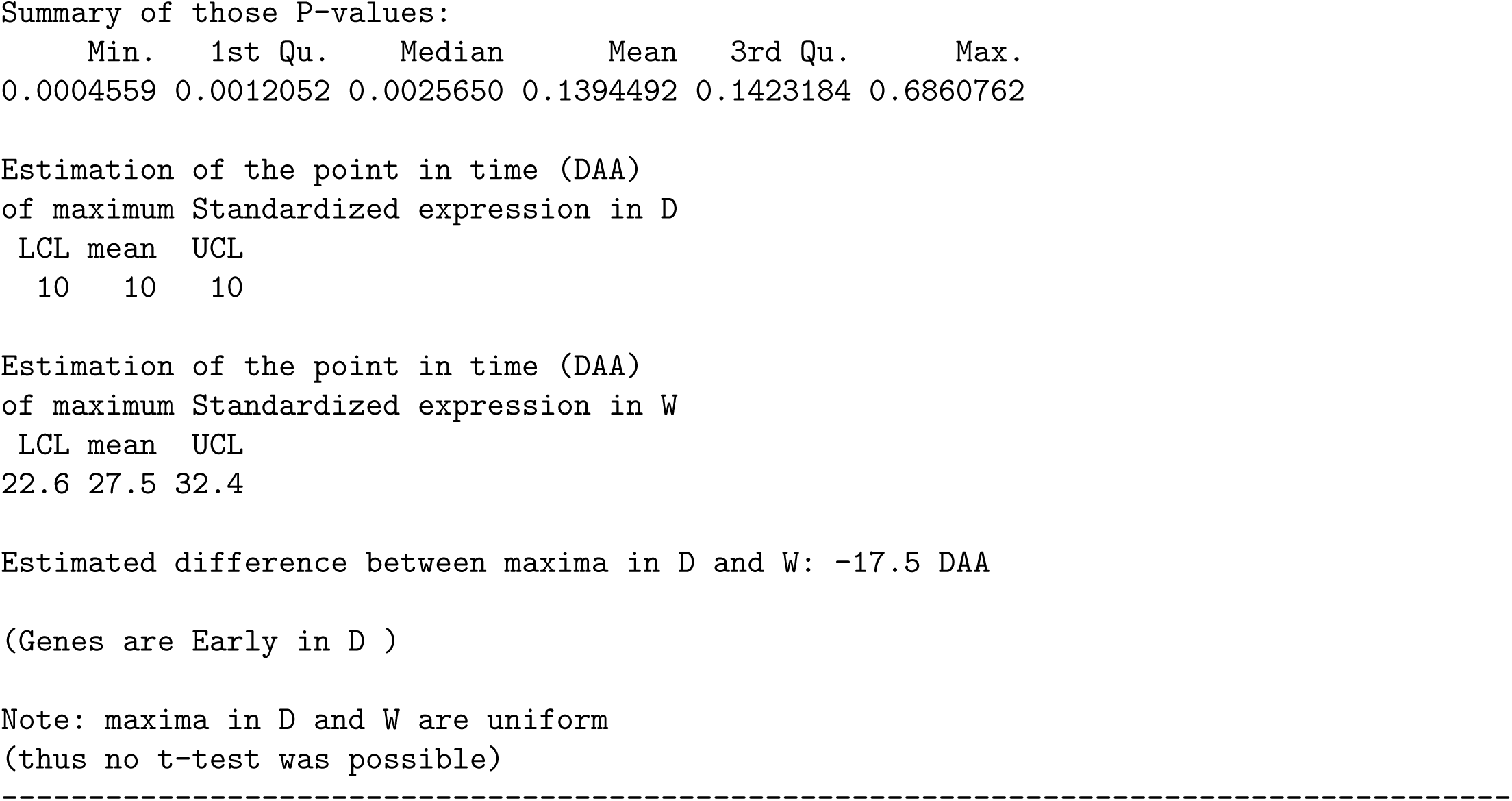

#### S-11.2. Analyses of GO biological process “Cell Cycle” having as target the D10W30 set of genes

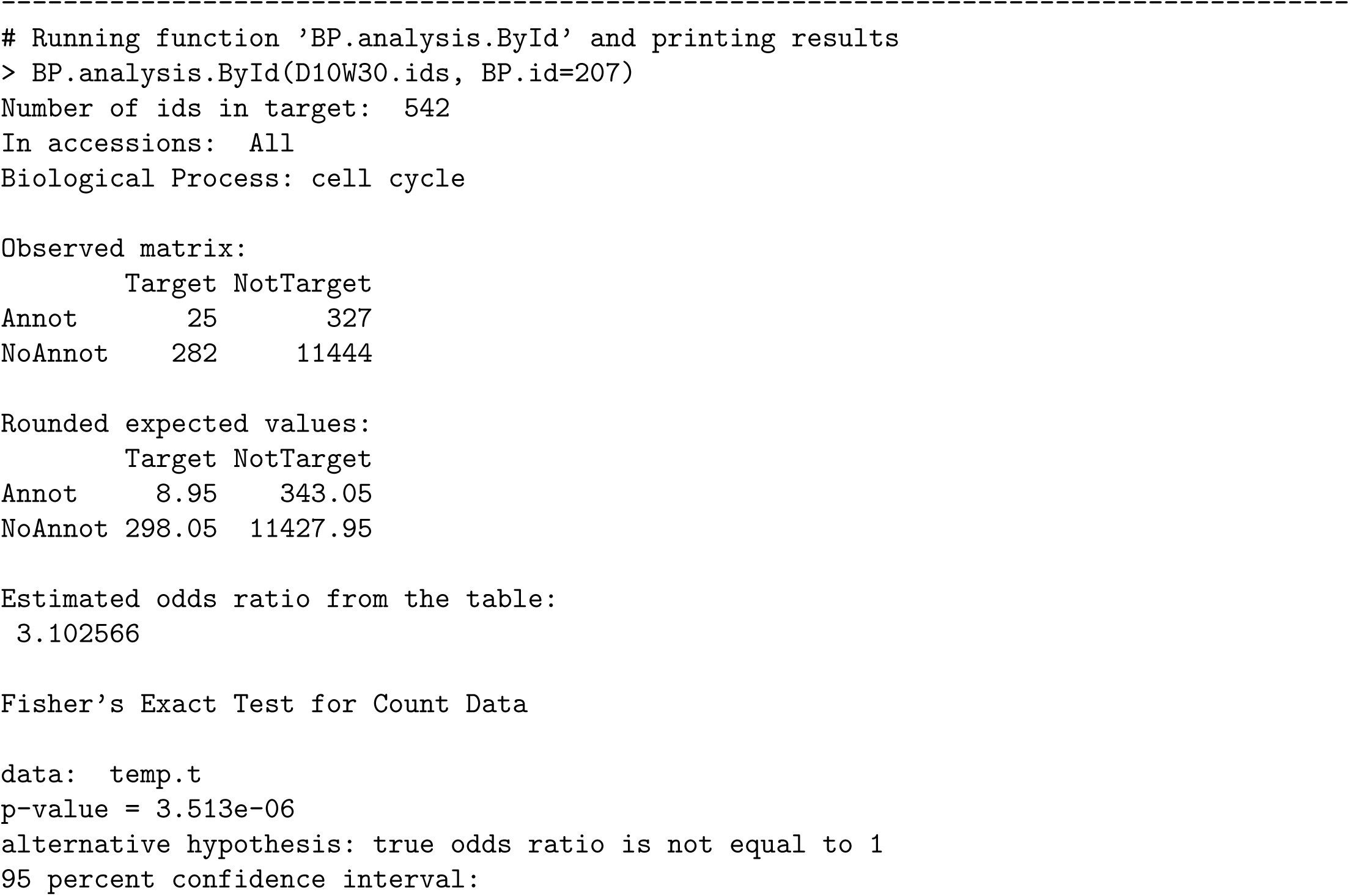
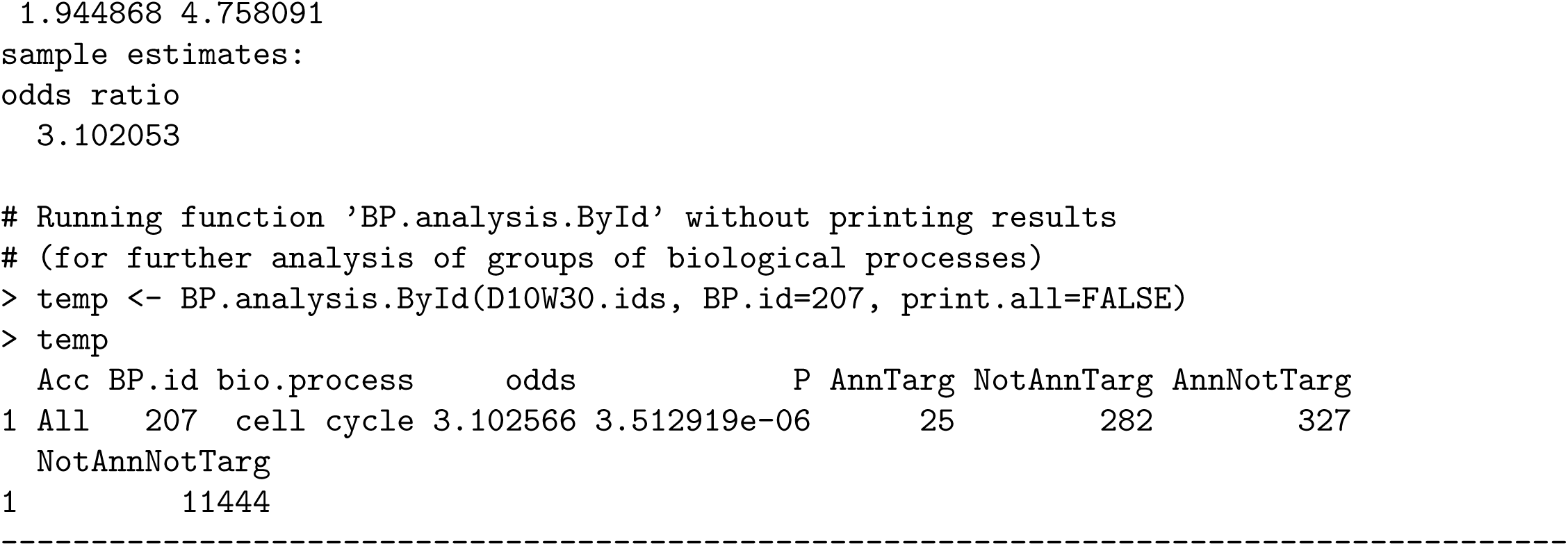

## Footnotes

1 FPKM stands for ‘number of Fragments Per Kilobase of transcript sequence per Millions base pairs sequenced’

